# Muscle Van Gogh-like 2 shapes the neuromuscular synapse by regulating MuSK signaling activity

**DOI:** 10.1101/2020.11.26.384925

**Authors:** Myriam Boëx, Julien Messéant, Steve Cottin, Marius Halliez, Stéphanie Bauché, Céline Buon, Nathalie Sans, Mireille Montcouquiol, Jordi Molgó, Muriel Amar, Arnaud Ferry, Mégane Lemaitre, Andrée Rouche, Dominique Langui, Asha Baskaran, Bertrand Fontaine, Laure Strochlic

## Abstract

The development of the neuromuscular junction (NMJ) requires dynamic trans-synaptic coordination orchestrated by secreted factors, including the morphogens of the Wnt family. Yet, how the signal of these synaptic cues is transduced, and particularly during the regulation of acetylcholine receptor (AChR) accumulation in the postsynaptic membrane remains unclear. We explored the function of Van Gogh-Like protein 2 (Vangl2), a core component of Wnt planar cell polarity signaling. We showed that the conditional genetic ablation of Vangl2 in muscle reproduces the NMJ differentiation defects in mice with constitutive Vangl2 deletion. These alterations persisted into adulthood with NMJs disassembly leading to an impairment of neurotransmission and motor function deficits. Mechanistically, we found that Vangl2 and the muscle-specific kinase MuSK acted in the same genetic pathway and that Vangl2 binds MuSK, thus controlling its signaling activity. Our results identify Vangl2 as a key player of the core complex of molecules shaping neuromuscular synapses and shed light on the molecular mechanisms underlying NMJ assembly.

## Introduction

The neuromuscular junction (NMJ) is a tripartite peripheral synaptic contact between presynaptic motor neurons, glial Schwann cells and postsynaptic striated skeletal muscle fibers. In vertebrates, this chemical synapse uses the excitatory neurotransmitter acetylcholine (ACh) to activate ACh receptors (AChRs) concentrated in the postsynaptic membrane, leading to rapid, reliable muscle contraction ^1,2^. NMJ formation requires a complex interplay of synaptic cues from several cellular compartments, triggering intracellular signaling cascades that control synaptic gene and molecular regulatory networks, thereby coordinating the precise alignment between pre- and postsynaptic domains. A disruption of this cellular polarization and *trans*-synaptic coordination leads to the progressive breakdown of synaptic connections, associated with disorders of neuromuscular transmission with major impacts on motor function. The receptor complex formed by MuSK (muscle specific kinase) and Lrp4 (LDL receptor-related protein 4), a member of the Low-Density-Related Protein family, constitutes the central scaffold orchestrating both NMJ formation and maintenance from the postsynaptic side ^3–9^. Until recently, the dogma has been that activation of this complex occurred exclusively *via* the nerve-derived anterograde factor Agrin ^6,9,10^. Agrin binds to the extracellular domain of Lrp4, stimulating the association of MuSK with Lrp4, and increasing MuSK phosphorylation and AChR clustering ^6,9–13^. However, our group and others overturned this view showing that signaling molecules controlling the early wiring of synaptic contacts, such as the Wnt secreted morphogens can mediate MuSK/Lrp4 activation ^3,14–17^. We notably showed that the binding of the MuSK cysteine-rich domain (CRD) to Wnts (i.e. Wnt4 and Wnt11) leads to the downstream activation of several Wnt signaling pathways. These include the canonical β-catenin-dependent, and the planar cell polarity (PCP) pathways, both required for key aspects of NMJ development, such as synapse-specific gene expression, AChR clustering and the retrograde control of motor axon outgrowth ^15,16,18^. However, despite the strong evidence of Wnt-elicited functions at the vertebrate NMJ ^1,19–22^, it remains unclear how Wnts transduce their signals, particularly in muscle, to drive NMJ formation and maintenance.

To get insight into this, we studied the function and molecular determinants of Van Gogh-Like protein 2 (Vangl2)-dependent Wnt/PCP signaling. Vangl2 is a highly conserved four pass transmembrane protein belonging to the core complex responsible for mediating Wnt/PCP signaling, which is known to cause cell polarity by modifying the cytoskeleton within the cell. Vangl2 is expressed in developing motor axons and is essential for axon outgrowth in the central nervous system (CNS) ^23–26^. In the rat CNS, Vangl2 localizes to the postsynaptic density, where it interacts with PSD95, suggesting a possible role in synaptogenesis and plasticity ^27,28^. We recently showed that Vangl2 is enriched in both nerve and muscle NMJ counterparts, that it interacts with Wnt4 and Wnt11 and that mouse embryos bearing the spontaneous *Vangl2* loop tail mutation, display strong defects of AChR clustering and a striking overgrowth of motor axons, similar to *MuSK*Δ*CRD* ^15^. This suggests that Vangl2-dependent processes contribute to presynaptic motor axon outgrowth and postsynaptic assembly. In this study, we generated transgenic mice with constitutive, muscle- and motoneuron-specific *Vangl2* deletions to demonstrate that the loss of Vangl2 function in muscle, but not in motoneurons phenocopies the NMJ alterations triggered by the constitutive ablation of *Vangl2* during embryonic development. These embryonic pre- and postsynaptic differentiation defects continued in adulthood as NMJs disassembled leading to the progressive failure of neuromuscular transmission and motor function deficits. *In vivo* genetic interaction experiments demonstrated that Vangl2 and MuSK belonged to the same signaling cascade, mediated partly by the binding of Wnt to MuSK. We showed that Vangl2 binds physically to MuSK and controls its signaling activity in muscle cells. A loss of Vangl2 function in the muscle inhibited RhoA activation and altered actin cytoskeletal organization leading to AChR clustering deficiency. Overall, our results identify Vangl2 as a new regulator of MuSK signaling playing a key role in postsynaptic assembly and reveal a previously uncharacterized function of Wnt/PCP in muscle relying on the Vangl2/MuSK interaction to shape mammalian neuromuscular connectivity.

## Methods

### Generation and genotyping of mouse lines

Mice were maintained in a controlled-temperature environment, under a 12-h light/12-h dark cycle, with *ad libitum* access to water and rodent chow diet. All experiments were performed in accordance with national and European guidelines for animal experimentation, with the approval of the institutional ethics committee (Institut du Cerveau et de la Moelle (ICM), N°A-75-1970). The *Vangl2*^*LoxP/LoxP*^ transgenic mouse, in which exon 4 of the *Vangl2* gene is flanked by LoxP sites, was described in detail by Ramsbottom et al. (2014). Cre driver mouse lines, including B6.C-Tg(CMV-Cre)1Cgn/J (CMV::Cre, Jax catalog no. 006054), B6.C-Tg(ACTA1-Cre)79Jme/J (HSA::Cre, Jax catalog no. 006149) and B6.129S1-*Mnx1*^*tm4*(*Cre*)*Tmj*^*/J* (HB9::Cre, Jax catalog no. 006600), were crossed with the *Vangl2*^*LoxP/LoxP*^ line to generate CMV::Cre;*Vangl2*^*LoxP/LoxP*^ (referred to hereafter as *Vangl2*^*-*/*-*^), HSA::Cre;*Vangl2*^*LoxP/LoxP*^ (*HSA-Vangl2*^*-*/*-*^) and HB9::Cre;*Vangl2*^*LoxP/LoxP*^ (*HB9-Vangl2*^*-*/*-*^) mice respectively. HSA-*Vangl2*^*LoxP/-*^; *MuSK*^+/*-*^ and HSA-*Vangl2*^*LoxP/-*^;MuSK^+/ΔCRD^ transheterozygous mice were obtained by crossing HSA::Cre;*Vangl2*^*LoxP/+*^ mice with *MuSK*^+/*-*4^ and *MuSK*^*+/*Δ*CRD* 18^ mice, respectively. *Vangl2*^*LoxP/LoxP*^ transgenic animals were genotyped by PCR on genomic DNA isolated from the tail, with the following primers: forward, 5’-ccgctggctttcctgctgctg-3’ and reverse, 5’-tcctcgccatcccaccctcg-3’ ^29^. The oligonucleotide primers used for CMV::Cre, HSA::Cre and HB9::Cre genotyping were as follows: CMV::Cre, forward, 5’-gcggtctggcagtaaaaactatc-3’ and reverse, 5’-gtgaaacagcattgctgtcactt-3’; HSA::Cre, forward, 5’-aagtgaagcctcgcttcc-3’ and reverse, 5’-cctcatcactcgttgcatcga-3’ ^30^; HB9::Cre, forward, 5’-tggctacccgtgatattgct-3’ and reverse, 5’-ccacagctcgctaggaggtgag-3’ ^30^.

### Antibodies

The following primary antibodies were used: rabbit polyclonal anti-MuSK (1:1,000; Merck Millipore, catalog no. ABS549), anti-HA.11 (1:100 for immunoprecipitation, Biolegend, catalog no. 902301), anti-HA (1:2500 for Western blotting, Abcam, catalog no. ab9110), anti-synaptophysin (Syn, 1:750, ThermoFisher Scientific, catalog no. 180130), anti-neurofilament (NF) 68kDa (1:500, Millipore, catalog no. AB9568), rat polyclonal anti-Vangl2 (clone 2G4, 1:200 for Western blotting, 1:400 for immunoprecipitation, 1:500 for immunostaining, Merck Millipore, catalog no. MABN750), mouse monoclonal IgG1k anti-GFP (clones 7.1 and 13.1, 1:100 for immunoprecipitation, Roche, catalog no. 11814460001), IgG2b anti-GFP (clone GT859, 1:5000 for Western blotting, Sigma, catalog no. SAB2702197), IgG1 anti-transferrin receptor (TfR, clone H68.4, 1:500, ThermoFisher Scientific, catalog no. 13-6800), IgG1 anti-GAPDH (1:10,000; Abcam, catalog no. ab8245). The following secondary antibodies for immunostaining were purchased from ThermoFisher Scientific: goat anti-rabbit IgG (H+L) Alexa Fluor Plus 594 conjugated (1:400, catalog no. A32740), goat anti-mouse IgG1 Alexa Fluor 488 conjugated (1:400, catalog no. A21121), goat anti-rabbit IgG (H+L) Alexa Fluor Plus 555 conjugated (1:400, catalog no. A32732). The horseradish peroxidase (HRP)- conjugated goat anti-rat IgG light chain specific (1:10,000; Jackson ImmunoReseach laboratories, catalog no. 112-035-175), anti-mouse IgG (H+L) (1:10,000; ThermoFisher Scientific, catalog no. 31430) and anti-rabbit IgG (H+L) (1:10,000; ThermoFisher Scientific, catalog no. 31460) secondary antibodies were used for Western blotting. The Alexa Fluor 488-conjugated phalloidin (1:400, catalog no. A12379), Alexa Fluor 488- and Alexa Fluor 594-conjugated α-bungarotoxin (α-BTX) (1:500, catalog no. B13422 and B13423) were purchased from ThermoFisher Scientific.

### Plasmids, shRNA constructs and lentivirus generation

The rat MuSK-HA cDNA plasmid has been described elsewhere ^16,18,31^. The mouse GFP-Vangl2 cDNA plasmid has been previously used ^15,32^. Short hairpin RNA (shRNA) constructs and lentiviral particles were generated at the vectorology core facility iVECTOR (ICM/CNRS, Paris, France). A shRNA against *Vangl2* mRNA was designed to knockdown the expression of *Vangl2*. ShVangl2 was directed against the *Vangl2* sequence 5’-ccggtgggagtcgtggagataaattcaagagatttatctccacgactcccacc-3’. As a negative control, we used a scrambled shRNA (shScr) directed against the sequence 5’-cccctcgtcatagcgtgcataggttcaagagacctatgcacgctatgacga-3’. Lentiviral particles were produced by co-expressing shVangl2 and shScr with the p8.91 encapsidation plasmid, and pHCMV-G encoding the vesicular stomatitis virus (VSV) glycoprotein-G in HEK293T cells with titration according to the iVECTOR protocol.

### Cell cultures, transfection and lentiviral infection

COS-7 cells (ATCC, catalog no. CRL-1651) and mouse C2C12 myoblasts (ATCC, catalog no. CRL-1772) were maintained in Dulbecco’s modified eagle medium (DMEM) GlutaMAX™ (Gibco-Life Technologies) supplemented with 10% or 20% (v/v) fetal bovine serum (Gibco-Life Technologies), respectively, and 1% (v/v) penicillin/streptomycin solution (Gibco-Life Technologies) at 37°C under an atmosphere containing 5% CO_2_. Cells were plated in six-well plates, at a density of 500 000 cells per well. COS-7 cells were transiently transfected with 2μg plasmid (MuSK-HA, or GFP-Vangl2 alone, or in combination) using Fugene® 6 reagent (Promega, catalog no. E2311), according to the manufacturer’s recommendations. ShVangl2 and shScr lentiviruses were used at a multiplicity of infection (MOI) of 100:1 for the transduction of C2C12 cells. Two days after lentivirus transduction, C2C12 cells were allowed to differentiate into myotubes in DMEM supplemented with 5% (v/v) horse serum (Gibco-Life Technologies) and 1% (v/v) penicillin/streptomycin solution, which were then harvested for downstream applications.

For primary muscle cell culture, hindlimb tibialis anterior (TA) muscles from E18.5 mouse embryos were explanted and digested in 0.2% collagenase type I (Gibco-Life Technologies, catalog no. 17100-017) in DMEM for 60 min at 37°C. Individual myofibers were dissociated with a Pasteur pipette coated with heat-inactivated 5% bovine serum albumin (Sigma). Pieces of muscle fiber were plated in growth medium consisting of DMEM GlutaMAX™ supplemented with 5% (v/v) horse serum, 0.5% (v/v) chicken embryonic extract (Seralab), 1% (v/v) penicillin streptomycin solution, 2% (v/v) fungizone and incubated for 4h at 37°C under an atmosphere containing 5% CO_2_. The supernatant was removed by centrifugation (16,000 x g, 20 min, 4°C) and the resulting pellet was resuspended in growth medium Matrigel-coated dishes (Corning, catalog no. 356231) for five days. Differentiation was induced by incubating primary myoblasts in DMEM GlutaMAX™ supplemented with 2% (v/v) horse serum, 1% (v/v) chicken embryonic extract, 1% (v/v) penicillin streptomycin solution, and 2% (v/v) fungizone.

### Duolink (®) *in situ* PLA technology

COS-7 cells were plated on coverslips and transfected with cDNA-containing plasmids as described above. Cells were fixed by incubating with 4% PFA in PBS for 15 min and then permeabilized by incubation with 0.5% Triton X-100 in PBS for 5 min at room temperature. They were blocked by incubating with Duolink blocking solution according to the manufacturer’s indications (Duolink^®^ *in situ* PLA kit, Sigma). Cells were incubated with rabbit polyclonal anti-HA.11 and mouse monoclonal IgG1k anti-GFP antibodies at 4°C overnight. Incubation with PLA probes, ligation, amplification and mounting were performed according to the Duolink kit instructions. The following PLA probes were used: anti-rabbit plus (Sigma, catalog no. DUO92002), and anti-mouse minus (Sigma, catalog no. DUO92004). Images were collected with an epifluorescence microscope equipped with an Apotome Slider module (ApoTome2, Zeiss), and collapsed into a single image. The PLA signals appeared as red fluorescent dots.

### Tissue protein extraction

Frozen extracts of muscle (diaphragm, *Tibialis anterior* (TA), gastrocnemius), brain and spinal cord were pulverized in liquid nitrogen and homogenized in 10 volumes of buffer containing 50 mM Tris, 10 mM EDTA, pH 8.3 and protease inhibitors (ThermoFisher Scientific, catalog no. 78438). An equal volume of buffer containing 0.125 M Tris, 4% SDS, 20% glycerol, pH 6.8 and protease inhibitors was added to the homogenates, which were then sonicated and incubated at 50°C for 20 min. Insoluble material was removed by centrifugation (16,000 x g, 20 min, 4°C). The concentration of protein in the supernatant was determined in a BCA assay (Pierce).

### Immunoprecipitation

Immunoprecipitation of transfected COS-7 cells or C2C12 myotubes transduced with shVangl2 or shScr was performed as described previously (Strochlic et al., 2012) with SureBeads™ protein G magnetic beads (BioRad, catalog no. 161-4023).

### Immunohistochemistry

Staining on whole-mount mouse embryo diaphragms and isolated adult muscle fibers was performed as previously described ^18^. For staining of primary myotubes, cells were fixed by incubation with 4% PFA in PBS for 15 min, permeabilized by incubation for 30 min with 0.5 Triton X-100 in PBS and blocked by incubation for 1 h at room temperature with 3% BSA, 5% goat serum and 0.5% Triton X-100 in PBS. Muscle cells were incubated overnight at 4°C with Alexa Fluor 488-conjugated phalloidin and Alexa Fluor TRITC-conjugated α-BTX. The cells were rinsed several times in PBS and were then flat-mounted in Vectashield® mounting medium (Vector Laboratories). For AChR cluster assays, myotubes were incubated overnight with, or without recombinant Wnt11 protein (10 ng/ml, R&D Systems, catalog no. 6179-WN/CF) and stained with Alexa Fluor 488-conjugated α-BTX in PBS for 1 h.

### Image acquisition and processing

All images were acquired with a Zeiss LSM-710 confocal laser scanning microscope equipped with a Plan-Apochromat 20 x / 0.8 NA objective, or Zeiss LSM-880 equipped with Plan-Apochromat 10 x/0.45 NA or 63 x/1.4 NA oil objectives. Z-stack images were acquired with LSM Zen Black software. The same laser power and parameter setting were used throughout, to ensure comparability. The confocal images presented are single-projected images obtained by overlaying sets of collected z-stacks.

A quantitative analysis of the NMJs on the embryonic left hemidiaphragm was performed, as previously detailed ^15^ with ImageJ software (version 2.0.0). The number and volume of AChR clusters were determined with the 3D object counter plugin-in ^33^ using the collected z-stacks. The number and length of both primary and secondary branches, defined as axons extending from the main phrenic nerve trunk, or the primary branches, respectively, were also quantified from maximum-intensity projections of the z-stacks. Morphometric analysis was performed on confocal z-stack projections of individual adult NMJs with an ImageJ-based workflow adapted from NMJ-morph ^34^. At least four hemidiaphragms or hundred isolated muscle fibers of each genotype were analyzed and quantified.

### Biotinylation of cell surface MusK

C2C12 myotubes transduced with shVangl2 or shScr were left untreated or were treated with recombinant human Wnt11 protein (10 ng/ml) for 5 min, 30 min, or 1 h at 37°C. Biotin-labelling of cell surface protein was performed as described previously ^18,35^.

### RhoA activation assay

RhoA activation was assessed with the G-LISA activation kit in accordance with the manufacturer’s instructions (Cytoskeleton Inc., catalog no. BK124-S). Active GTP-bound RhoA was detected with HRP-conjugated anti-RhoA antibodies. The HRP signal was quantified by measuring absorbance at 490 nm with a microplate spectrophotometer (Spark^®^, Tecan).

### Electron microscopy

Ultrastructural analysis was performed as previously described ^18^, with minor modifications. Samples were incubated twice in acetone for 10 min, overnight in 50%–50% Epon-50% acetone at 4°C in airtight closed container, then embedded in Epon and incubated 48h at 60°C for polymerization. The 70 nm sections were cut with an ultramicrotome (ultracut UC7, Leica) and collected on copper grids. Sections were contrasted for 10 min with 2% uranyl acetate and 4 min with Reynold’s lead citrate at room temperature. The ultra-thin sections were visualized with a CM120 Biotwin Philips transmission electron microscope operating at 80 kV equipped with a high-resolution wide-angle camera Morada SIS managed by Item software.

### *In vitro* isometric tension analyses of diaphragm muscles

We analyzed the contractile properties of diaphragm muscles *in vitro* on isolated left phrenic nerve-hemidiaphragm preparations, as previously described ^18,36^. In brief, P180 adult mice were anesthetized with isoflurane and killed by cervical dislocation. The left phrenic nerve-hemidiaphragm preparation was mounted in a silicone-lined bath superfused with an oxygenated normal Krebs-Ringer solution. A suction microelectrode allowed phrenic-nerve stimulation with current pulses (0.15 ms duration) at frequencies indicated in figures. An electrode-assembly placed along the length of the hemidiaphragm allowed direct muscle stimulation. Signals from the force sensor were amplified and digitized with a computer equipped with an analog-to-digital interface board (Digidata 1550B with pCLAMP 10.7 software, Molecular Devices). All experiments were performed at 22 ± 0.5°C.

### *In situ* isometric force analysis and electroneuromyography of hindlimb muscles

*In situ* isometric contraction of the TA muscle was achieved, as previously described ^37^. A blind analysis was performed by the investigator. Signals from the force sensor were amplified and digitized with a computer equipped with an analog-to-digital interface board (BioAmp, ADInstruments) with LabChart 8 software (ADInstruments). For electroneuromyography, two monopolar reference and recording electrodes were inserted, one close to the patella tendon, and the other in the middle of the TA muscle. A monopolar ground electrode was also inserted into the contralateral hindlimb muscle. The sciatic nerve was stimulated with series of 10 stimuli at 1, 5, 10, 20, 30 and 40 Hz. Compound muscle action potentials (CMAPs) were amplified (BioAmp, ADInstruments), acquired with a sampling rate of 100 kHz, filtered with a 5 kHz low-pass and a 1 Hz high-pass filter (Powerlab 4/25, ADInstruments), and peak-to-peak amplitudes were analyzed with LabChart 8 software (ADInstruments). During all experiments mouse body temperature was maintained at 37°C with radiant heat. After recordings, the mice were killed by cervical dislocation and the muscles were collected.

### Statistical analysis

All statistical analyses and graphs were plotted with GraphPad Prism 8.0 software. Data are presented as means ± SEM. Mann-Whitney U-tests were used for comparisons of two groups. For comparisons of more than two groups simultaneously, one- or two-way ANOVA was performed, followed by an appropriate multiple comparison test (as detailed in figure legends). Values of *p* ≤ 0.05 were considered statistically significant. Each experiment was conducted of at least three times.

## Results

### Constitutive Vangl2 deletion impairs embryonic NMJ formation

We investigated the specific function of Vangl2 during NMJ formation and overcame the putative dominant negative effect of the Vangl2 loop tail mutation ^38,39^, by studying Vangl2-deficient mice (*Vangl2*^*-*/*-*^) generated by crossing *Vangl2*^*LoxP/LoxP*^ transgenic mice, in which exon 4 of *Vangl2* is flanked by LoxP sites, with mice expressing the Cre gene under the control of the ubiquitous cytomegalovirus (CMV) promoter **(Fig. 1A)**. Homozygosity for Vangl2 deletion causes a severe neural tube closure defect lethal at birth, a hallmark of impaired PCP signaling, consistent with the described role of Vangl2 during mouse development ^40,41^. Genotyping and Western blot analyses of Vangl2 expression in several tissue extracts, including brain, spinal cord and muscles, from control and *Vangl2*^*-*/*-*^ embryos confirmed the constitutive deletion of Vangl2 (**Fig. 1B and 1C**). An analysis of the morphological phenotype of NMJs was performed by staining left hemidiaphragms from E18.5 control and *Vangl2*^*-*/*-*^ embryos with anti-neurofilament (NF) and anti-synaptophysin (Syn) antibodies and α-bungarotoxin (α-BTX) to label the phrenic nerve, terminal branching and acetylcholine receptors (AChR), respectively **(Fig. 1D)**. In *Vangl2*^*-*/*-*^ embryos, as in control, the main nerve is located in the central region of the diaphragm, and oriented perpendicular to the muscle fibers (**Fig. 1D**). Nerve terminals reach and innervate AChR clusters on muscle fibers adjacent to the nerve trunk. Vangl2 mutant mice presented pre- and postsynaptic NMJ differentiation defects (**Fig. 1D)**. The number (control, 100 ± 3.8%; *Vangl2*^*-*/*-*^, 73 ± 3.4%; *p* < 0.001) and the volume (control, 100 ± 2.4 μm^3^; *Vangl2*^*-*/*-*^, 72 ± 2.3 μm^3^; *p* < 0.001) of AChR clusters were ∼30% lower in *Vangl2*^*-*/*-*^ embryos to than in control (**Fig. 1E-F**). Moreover, primary and secondary nerve branches were 45% (control, 67 ± 4.9 μm; *Vangl2*^*-*/*-*^, 97 ± 8.5 μm; *p* < 0.01) and 150% (control, 85 ± 4.8 μm; *Vangl2*^*-*/*-*^, 213 ± 8.9 μm; *p* < 0.001) longer, respectively, in *Vangl2*^*-*/*-*^ embryos to than in control **(Fig. 1G-H)**. No difference in the number of primary and secondary nerve branches was detected (**Fig. 1I-J**). Interestingly, the number of aneural AChR clusters was also found to be smaller in E13.5 *Vangl2*^*-*/*-*^ embryos than in controls with no effect on presynaptic nerve localization and growth (**supplemental Fig. 1A-C**), suggesting that Vangl2 is required from the first step of NMJ formation.

**Figure 1.**
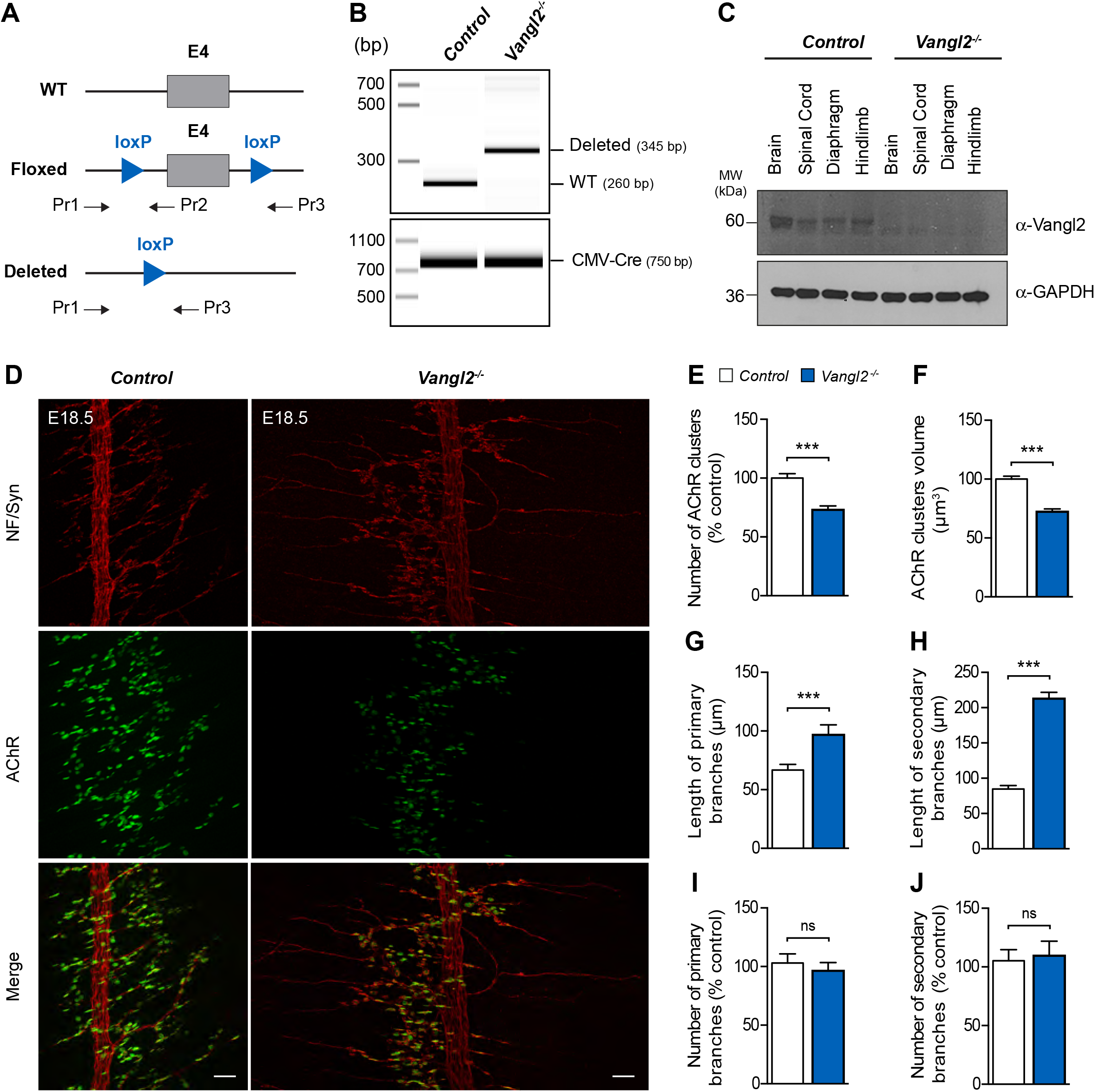
Loss of Vangl2 function impairs NMJ formation. **A**. Diagrams of the control and floxed alleles of the mouse *Vangl2* gene. Pr1, Pr2 and Pr3 correspond to genotyping primers. E4, exon 4. **B**. Genotyping analysis of control and *Vangl2*^*-/-*^ mouse embryos. DNA was extracted from the tail and subjected to PCR analysis. Left, molecular weight (bp). Cre primers generate 750 bp products. Vangl2^LoxP^ primers generate 260 bp products for the control allele and 345 bp products for the deleted allele. **C**. Western-blot analysis of Vangl2 levels in homogenates obtained from various tissues (brain, spinal cord, and skeletal muscles, diaphragm and hindlimb tissues) from E18.5 control and *Vangl2*^*-*/*-*^ mouse embryos. GAPDH, loading control. MW, molecular weight (kDa). **D**. Representative confocal images of whole-mount left hemidiaphragms from E18.5 control and *Vangl2*^*-*/*-*^ mouse embryos stained with α-BTX (green, RACh), anti-neurofilament (red, NF, phrenic nerve) and anti-synaptophysin (red, Syn, nerve terminals) antibodies. Scale bars in the merged images, 50 μm. **E-J**. Quantitative analysis of AChR cluster numbers (E), volume (F), length of primary (G) and secondary (H) nerve branches, as well as number of primary (I) and secondary (J) nerve branches. Data are means ± SEM, ns, non-significant; \*\*\**p* < *0.001, N* = *4* embryos per genotype, Mann-Whitney *U* test.

### Aberrant NMJ formation in the absence of muscle Vangl2

Since Vangl2 is present in both the muscle postsynaptic membrane, and in presynaptic nerve terminals ^15^, we generated conditional mutants to discriminate the role of Vangl2 in both tissues using promoters typically used to drive expression of the Cre recombinase in the myotomal regions of somites (HSA) or in motoneurons (HB9) ^42–45^. A genotyping analysis of HSA::Cre; *Vangl2*^*loxP/loxP*^ (*HSA-Vangl2*^*-*/*-*^) and HB9::Cre; *Vangl2*^*loxP/loxP*^ (*HB9-Vangl2*^*-*/*-*^) mutants showed the deletion of the *Vangl2* allele in DNA isolated from skeletal muscle or spinal cord, respectively, confirming the tissue-specific deletion (**Fig. 2A**). A Western blot also showed a strong decrease of Vangl2 protein levels in the corresponding specific tissue homogenates of *HSA-Vangl2*^*-*/*-*^ and *HB9-Vangl2*^*-*/*-*^ mice compared to control (**Fig. 2B**). *HB9-Vangl2*^*-*/*-*^ embryos were phenotypically indistinguishable from controls, whereas *HSA-Vangl2*^*-/-*^embryos displayed abnormal NMJ differentiation **(Fig. 2C-I)**. There were 30% fewer AChR clusters (*LoxP/+*, 100 ± 3.6%; *HSA*-*Vangl2*^*-*/*-*^, 71 ± 6%; *p* < 0.01; **Fig. 2D**), and these clusters occupied a volume 53% smaller (*LoxP/+*, 226 ± 15.7 μm^3^; *HSA*-*Vangl2*^*-*/*-*^, 105 ± 7.8 μm^3^; *p* < 0.01; **Fig. 2E**) than in control embryos. Moreover, *HSA-Vangl2*^*-*/*-*^ embryos displayed extension of motor axons, bypassing the remaining AChR clusters and growing aberrantly toward the muscle periphery. The primary and secondary nerve branches were 38% (*LoxP/+*, 41 ± 2.4 μm; *HSA*-*Vangl2*^*-*/*-*^, 78 ± 6.4 μm; *p* < 0.001) and 90% (*LoxP/+*, 77 ± 4 μm; *HSA*-*Vangl2*^*-*/*-*^, 122 ± 7μm; *p* < 0.001) longer, respectively, but their number remained unchanged (**Fig. 2F-I**). Overall, these findings indicate that Vangl2 expression is required for NMJ differentiation in muscle, but dispensable in motoneurons.

**Figure 2.**
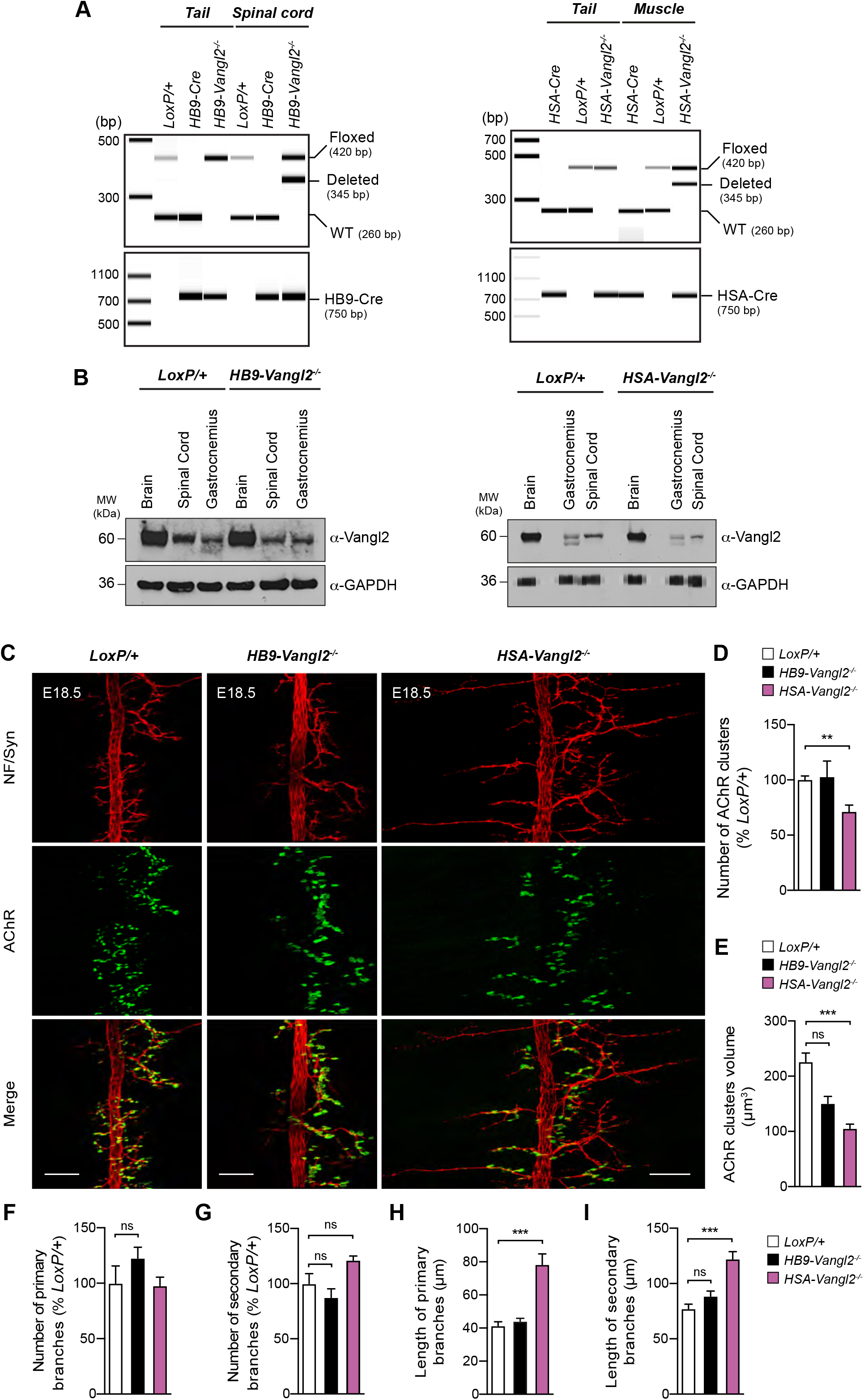
Muscle Vangl2 is involved in NMJ differentiation whereas motoneuron Vangl2 is not. **A**. Genotyping analysis of *Vangl2*^*LoxP/+*^ (*LoxP/+*, control), *HB9-Vangl2*^*-*/*-*^ and *HSA-Vangl2*^*-*/*-*^ mouse embryos. DNA was extracted from the tail, hind-limb muscles or spinal cord and subjected to PCR analysis. Left, molecular weight (bp). Cre primers generate 750 bp products. Vangl2^LoxP^ primers generate 260 bp products for the WT allele, 345 bp products for the deleted allele and 420 bp products for the floxed allele. The *HSA::Cre;Vangl2*^*LoxP/-*^ (*HSA-Vangl2*^*-*/*-*^) and *HB9::Cre;Vangl2*^*LoxP/-*^ (*HB9-Vangl2*^*-*/*-*^) genotypes were considered as conditional knockouts. **B**. Western-blot analysis of Vangl2 levels in homogenates from various tissues (brain, gastrocnemius muscle, and spinal cord tissues) from E18.5 *LoxP/+, HSA-Vangl2*^*-*/*-*^ and *HB9-Vangl2*^*-*/*-*^ mouse embryos. GAPDH, loading control. MW, molecular weight (kDa). **C**. Representative confocal images of whole-mount left-hemidiaphragms from E18.5 *Vangl2*^*LoxP/+*^ (*LoxP/+*, control), *HB9-Vangl2*^*-*/*-*^ and *HSA-Vangl2*^*-*/*-*^ mouse embryos stained as in **Fig. 1**. Scale bars in the merged images, 50 μm. **D-I**. Quantitative analysis of AChR cluster numbers (D), volume (E), number of primary (F) and secondary (G) nerve branches, and the length of primary (H) and secondary (I) nerve branches. Data are means ± SEM, ns, non-significant; \*\**p* < *0.01;* \*\*\**p* < *0.001, N* = *6* embryos per genotype, one-way ANOVA, Tukey’s test.

### Conditional Vangl2 deletion in muscle affects NMJ architecture

To evaluate the consequences of an early Vangl2 deletion in muscle, we first analyzed the NMJ morphology of isolated *tibialis anterior* (TA) muscle fibers from P180 mutant mice, comparing our findings with those of control TA (**Fig. 3A-G**). The NMJs of *HSA-Vangl2*^*-*/*-*^ mice adopted a characteristic ‘pretzel-like’ shape, but postsynaptic structures were discontinuous, with isolated AChR clusters. Quantitative analyses showed that the AChR clusters were smaller in *HSA-Vangl2*^*-*/*-*^ mice than in the control (*LoxP/+*, 641 ± 24.7 μm^2^; *HSA-Vangl2*^*-*/*-*^, 573 ± 18.9 μm^2^; *p* < 0.05) (**Fig. 3B**). *HSA-Vangl2*^*-*/*-*^ mice had significantly less compact NMJs, confirming a dispersion of the AChR clusters (**Fig. 3C**). We next determined the index of AChR fragmentation by measuring the number of continuous AChR-stained structures per synapse. The mean level of AChR fragmentation was higher in *HSA-Vangl2*^*-*/*-*^ mice than in control (*LoxP/+*, 1.0 ± 0.07; *HSA-Vangl2*^*-*/*-*^, 1.6 ± 0.07; *p* < 0.001) (**Fig. 3D**). None of the mutant NMJs analyzed had a perfect contiguous postsynaptic shape (i.e. number of AChR fragments = 1), and the number of NMJs with more than five AChR fragments was 2.7 times higher in *HSA-Vangl2*^*-*/*-*^ than in control (**Fig. 3E**). Despite these deficits, nerve terminals were juxtaposed to the postsynaptic AChRs and the area of the nerve terminals and the overlap between pre- and postsynaptic counterparts (i.e. AChR area covered by nerve terminals) were similar in *HSA-Vangl2*^*-*/*-*^ mice and control (**Fig. 3F-G**). We further analyzed the morphological alterations to muscle and NMJs at the ultrastructural level, by electron microscopy on TA muscles from P180 *HSA-Vangl2*^*-*/*-*^ and *LoxP/+* mice. *HSA-Vangl2*^*-*/*-*^ mice had fewer, more highly disorganized junctional folds than control (**Fig. 3H-K**) and a subset of mutant muscle specimens had disrupted sarcomeric regions (#, **Fig. 3I**). We also observed some NMJs with motor axon withdrawal, consistent with a process of muscle denervation (*, **Fig. 3K**).

**Figure 3.**
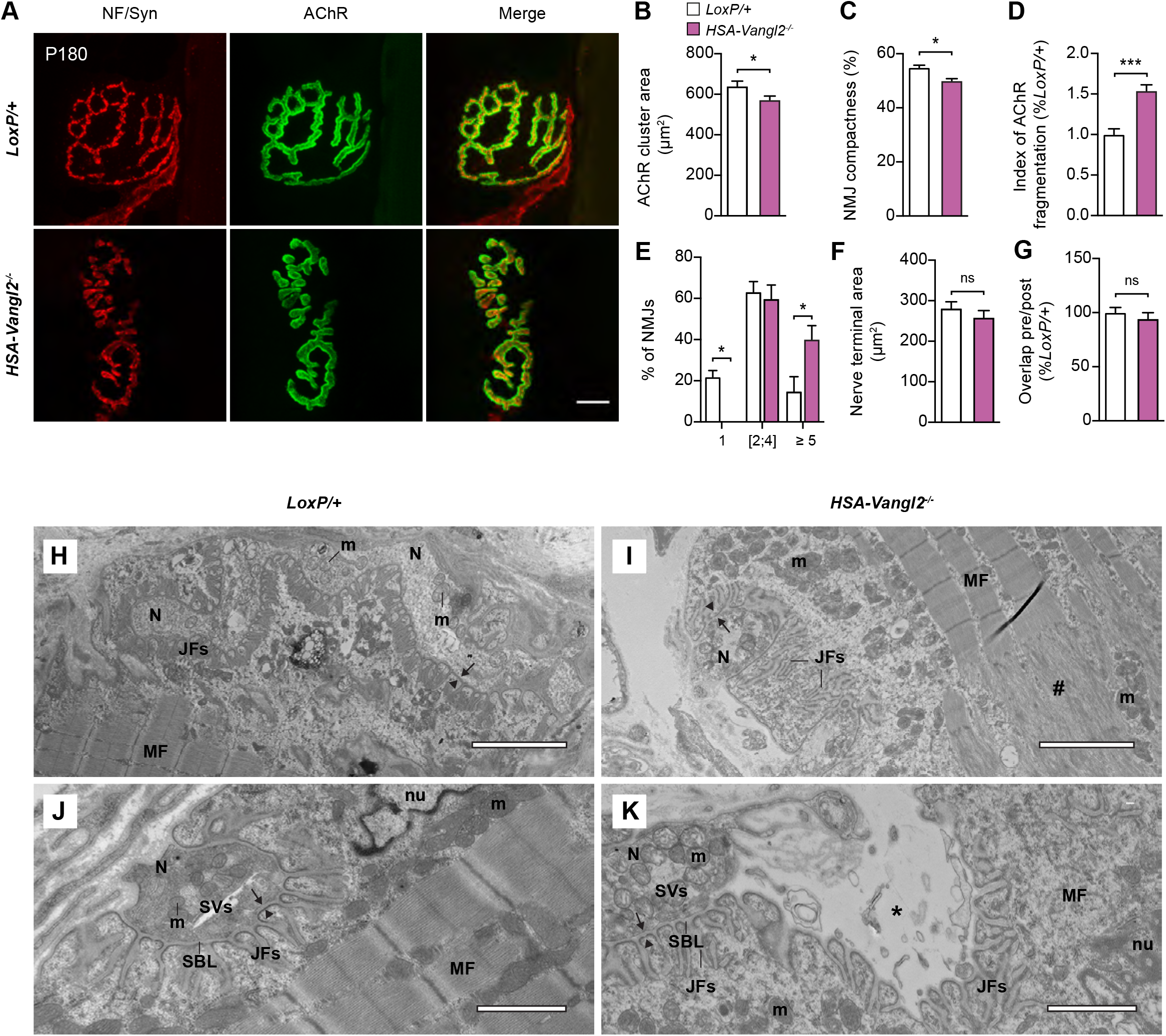
Abnormal NMJ structure and ultrastructure in adult *HSA-Vangl2*^*-*/*-*^ mice. **A**. Representative confocal images of isolated muscle fibers from P180 *LoxP/+* and *HSA-Vangl2*^*-*/*-*^ TA muscle stained with α-BTX (green, RACh), anti-NF and anti-Syn antibodies (red). Scale bar in the merged image, 10 μm. **B-G**. Quantitative analysis of **A**. The percentage of NMJs with an unfragmented postsynaptic apparatus or with at least 2 discontinuous AChR fragments is represented in **E**. Data are means ± SEM. ns, non-significant, \**p* < *0.05*, \*\*\**p* < *0.01, N* = *6* mice per genotype, Mann-Whitney U test (B-D, F-G) or two-way ANOVA, Sidak’s test (E). **H-K**. Representative electron micrographs of NMJs from P180 *LoxP/+* and *HSA-Vangl2*^*-*/*-*^ TA. N, nerve; SVs, synaptic vesicles; MF, muscle fiber; SBL, synaptic basal lamina; JFs, junctional folds; m, mitochondrion; nu, nucleus. Black arrows, presynaptic membrane; black arrowheads, postsynaptic membrane. *, axon withdrawal. #, disorganized sarcomeres. Scale bars, 3 μm in H-I; 1.25 μm in J-K, *N* = *3* mice per genotype.

### Altered synaptic transmission in mice with conditional Vangl2 deletion in muscle

We then investigated whether these changes in NMJ morphology resulted in motor function deficits. We first measured the nerve-evoked activity of TA muscles from P180 *HSA-Vangl2*^*-*/*-*^ mice and control *LoxP/+*, by assessing the maximal force produced *in situ* in response to tetanic sciatic nerve stimulation (**Fig. 4A-C**). Both the absolute force and specific force were ∼35% lower in *HSA-Vangl2*^*-*/*-*^ than in control muscle (absolute force: *LoxP/+*, 978 ± 30.3 mN; *HSA-Vangl2*^*-*/*-*^, 648 ± 27.4 mN; specific force: *LoxP/+*, 18 ± 1.1 mN/mg; *HSA-Vangl2*^*-*/*-*^, 12 ± 0.7 mN/mg; *p* < 0.05). To determine whether this muscle weakness resulted from a neuromuscular transmission failure, we recorded the compound muscle action potentials (CMAPs) triggered by 10 repetitive submaximal stimuli (**Fig. 4D-F**). The sciatic nerve was stimulated at different frequencies (1, 5, 10, 20, 30 and 40 Hz) and the CMAP amplitudes of *HSA-Vangl2*^*-*/*-*^ and *LoxP/+* TA muscles were recorded peak-to-peak. CMAP amplitudes between the first and 10^th^ stimuli were similar with controls, regardless stimulation frequency (**Fig. 4D-E**). However, CMAP amplitude at the 10^th^ stimulus was significantly smaller than at the first stimulus, for stimulation frequencies of 20 Hz or greater in *HSA-Vangl2*^*-*/*-*^ mice (**Fig. 4E**). At 40 Hz, for example, a decrease in CMAP amplitude was detected from the fifth stimulus onwards in *HSA-Vangl2*^*-*/*-*^ mice, resulting in a decrement of ∼13% (**Fig. 4F**). Thus, a loss of Vangl2 function in muscle contributes to muscle weakness due to a progressive loss of neurotransmission efficiency.

**Figure 4.**
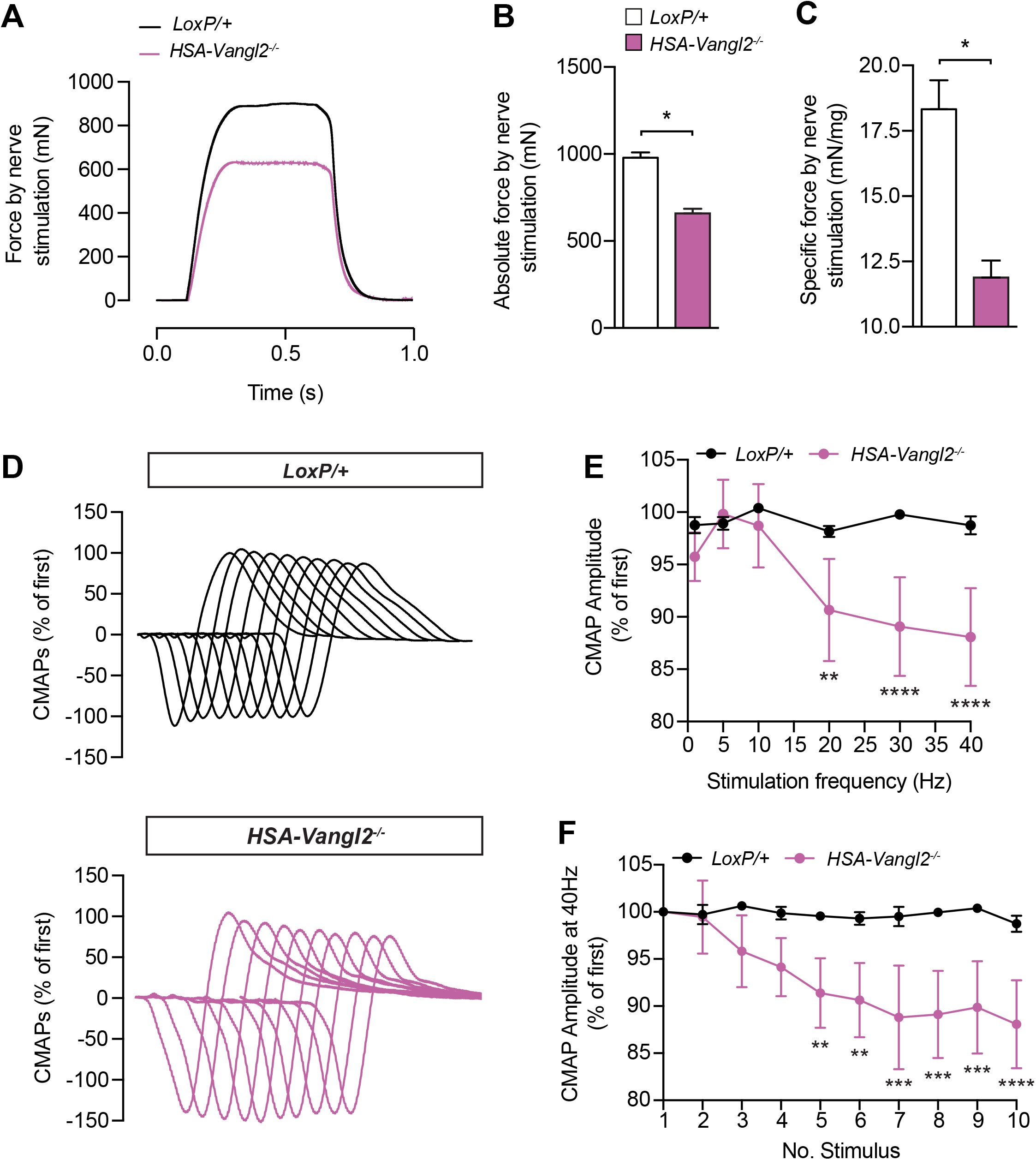
Reduced nerve-evoked tetanic force and CMAP in the tibialis anterior muscle of adult *HSA-Vangl2*^*-*/*-*^ mice. **A**. Representative tetanic contractions evoked by sciatic nerve stimulation at 100 Hz in *LoxP/+* and *HSA-Vangl2*^*-*/*-*^ P180 TA muscle. **B-C**. Quantitative analysis of nerve-evoked absolute maximal (B) and specific (C) muscle forces. Data are means ± SEM, ns, non-significant; \**p* < *0.05, N* = *4* mice per genotype, Mann-Whitney *U* test. **D**. Representative CMAP curves for P180 *LoxP/+* and *HSA-Vangl2*^*-*/*-*^ TA in response to ten repetitive sciatic nerve stimulations at 40 Hz. Curves are shown in a stack succession. **E**. Mean amplitude of the 10th CMAP, expressed as percentage of the first, at different nerve stimulation frequencies (1, 5, 10, 20, 30 and 40 Hz). **F**. Amplitude of 10 CMAPs, expressed as percentage of the first, at 40 Hz. Data are means ± SEM, ns, non-significant; ** *p*< *0.01*, \*\*\**p* < *0.001*, \*\*\*\**p* < *0.0001, N* = *4* mice per genotype, two-way ANOVA, Dunnet’s test.

Interestingly, mutant diaphragm muscles displayed a similar neurotransmission defect. Twitches and tetanic forces evoked by direct (muscle) or indirect (phrenic nerve) electrical stimulation at different frequencies (20, 40, 60 and 80 Hz) were measured *in vitro* on left phrenic nerve/hemidiaphragm preparations from P180 *LoxP/+* and *HSA-Vangl2*^*-*/*-*^ mice (**Fig. 5**). The contraction elicited by single or tetanic muscle stimulation did not differ significantly between *HSA-Vangl2*^*-*/*-*^ and *LoxP/+* mice (**Fig. 5A-C; Table 1**). However, the amplitude of the nerve-evoked muscle force was significantly smaller in hemidiaphragms from *HSA-Vangl2*^*-*/*-*^ mice than in control mice, in response to both single and tetanic phrenic nerve stimulation (decrease in the strength of muscle twitch: single twitch, 52%; tetanic stimuli, 20 Hz, 56%; 40 Hz, 56%; 60 Hz, 52%; 80 Hz, 38%) (**Fig. 5D-F; Table 1**). Similar results were obtained when twitch or tetanic forces were normalized by muscle weight (**Table 1**). These results highlight a diaphragm weakness in *HSA-Vangl2*^*-*/*-*^ mice caused by a neuromuscular transmission deficit.

**Figure 5.**
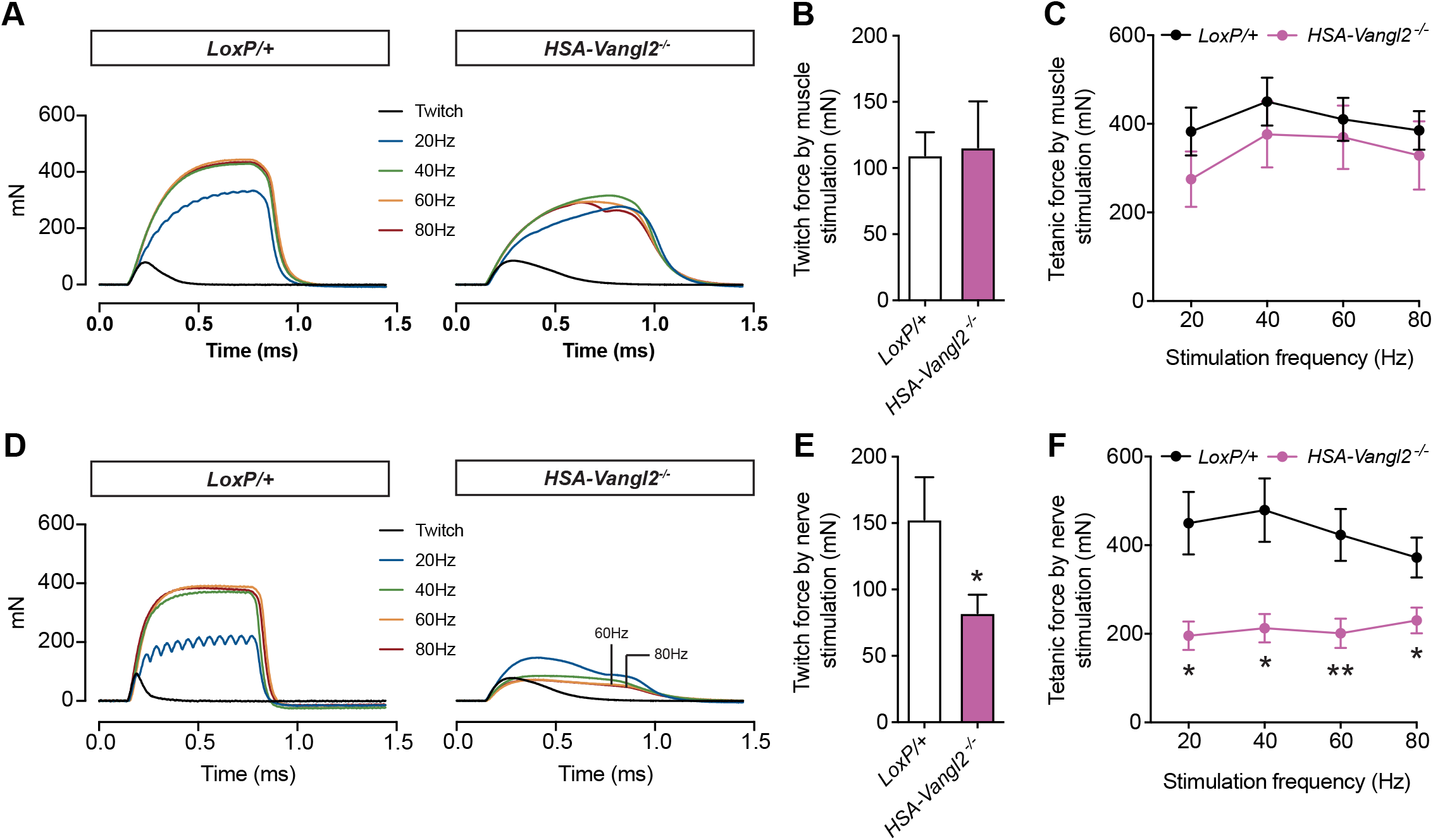
Decreased contraction following nerve stimulation of the isolated diaphragm from adult *HSA-Vangl2*^*-*/*-*^ mice. **A**. Representative twitch and tetanic contractions evoked by direct muscle stimulation in isolated P180 *LoxP/+* and *HSA-Vangl2*^*-*/*-*^ hemidiaphragms. The muscle was stimulated with single or tetanic stimuli at 20, 40, 60 and 80 Hz. **B-C**. Quantitative analysis of twitch (B) or tetanic forces (C) evoked by muscle stimulation. **D**. Representative twitch and tetanic contractions evoked by phrenic nerve stimulation in isolated P180 *LoxP/+* and *HSA-Vangl2*^*-*/*-*^ hemidiaphragms. The phrenic nerve was stimulated with single or tetanic stimuli at 20, 40, 60 and 80Hz. **E-F**. Quantitative analysis of twitch (E) or tetanic forces (F) evoked by phrenic nerve stimulation. Data are means ± SEM. \**p* < *0.05*, \*\**p* < *0.01, N* = *9* mice per genotype, Mann-Whitney *U* test (B, E) or two-way ANOVA, Dunnet’s test (C, F).

### Muscle Vangl2 cooperates with MuSK to orchestrate the NMJ formation

We investigated the molecular mechanisms by which muscle Vangl2 regulates NMJ formation, by testing the hypothesis that MuSK and Vangl2 act via the same signaling pathway required for postsynaptic differentiation. Given that extracellular Wnt4 and Wnt11 interact with MuSK *via* its Frizzled-like cysteine-rich domain (CRD), but also interact with Vangl2 ^15–17^, we first tested whether Vangl2 physically bound MuSK (**Fig. 6A-B**). We cotransfected COS-7 cells with plasmids encoding MuSK-HA and GFP-Vangl2 and performed reciprocal co-immunoprecipitation with anti-HA and anti-GFP antibodies. MuSK-HA co-immunoprecipitated with GFP-Vangl2 and *vice-versa* (**Fig. 6A**). We confirmed this Vangl2/MuSK interaction in a proximity ligation assays (PLA). Positive PLA signals (red spots) were detected in COS-7 cells co-expressing MuSK-HA and GFP-Vangl2 confirming the interaction (**Fig. 6B**).

**Figure 6.**
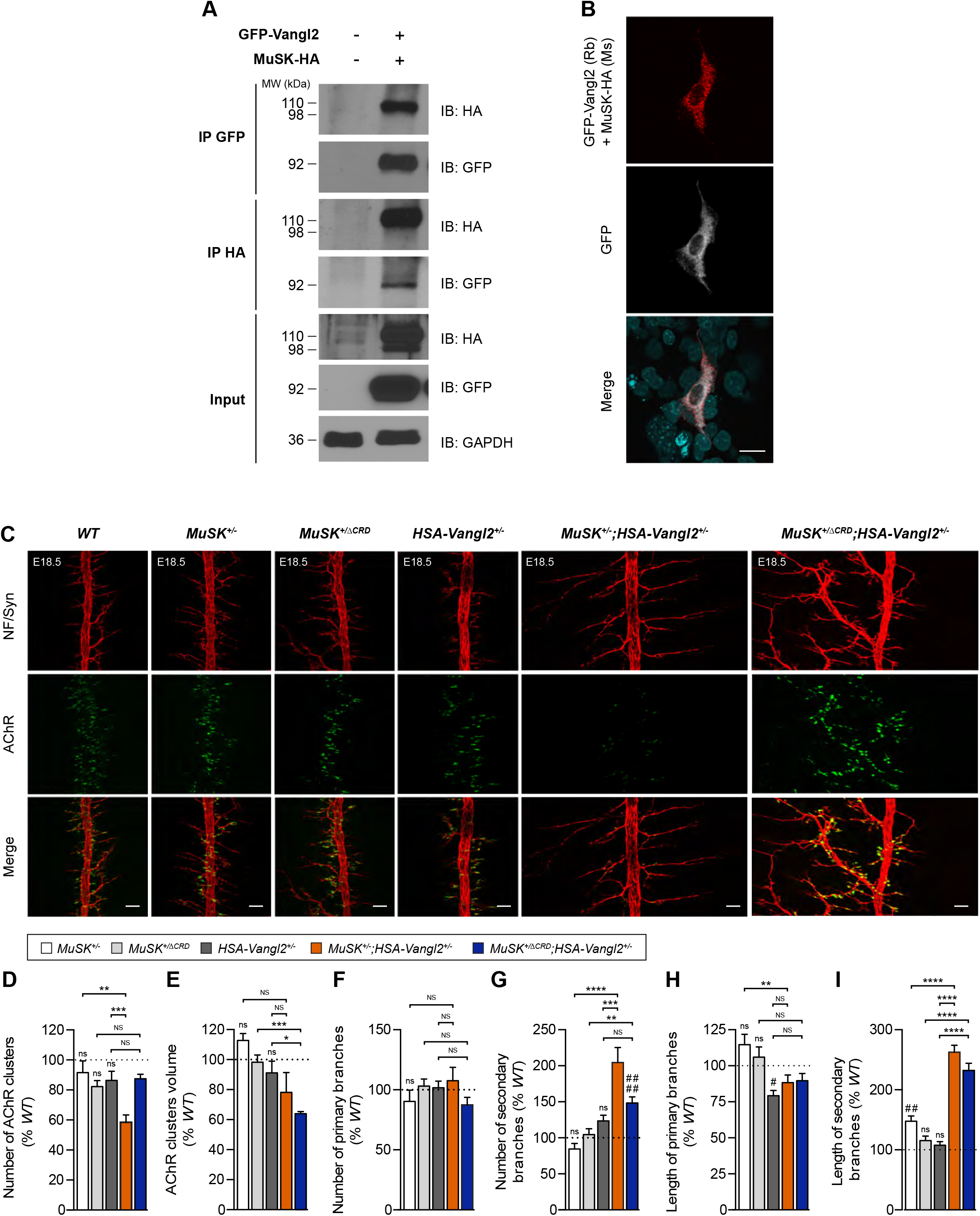
Genetic and physical interaction between Vangl2 and MuSK. **A**. Co-immunoprecipitation of GFP-Vangl2 and MuSK-HA in COS-7 cells. Western blots of cell lysates (Input) reveal the presence of the tagged proteins. GAPDH, loading control. MW, molecular weight (kDa). IP, immunoprecipitation. IB, immunoblotting. **B**. Duolink proximity ligation assay (PLA) in COS-7 cells expressing GFP-Vangl2 and MuSK-HA performed with anti-GFP (rabbit, Rb) and anti-HA (mouse, Ms) antibodies. PLA signal, red fluorescent dots. Blue in merged image, nuclei. Scale bar in the merged image, 50 μm. **C**. Representative confocal images of whole-mount left hemidiaphragms from E18.5 control, *MuSK*^+/*-*^, *MuSK*^*+/*Δ*CRD*^, *HSA-Vangl2*^+/*-*^, *MuSK*^+/*-*^;*HSA-Vangl2*^+/*-*^, *MuSK*^+/Δ*CRD*^;*HSA-Vangl2*^+/*-*^ mouse embryos stained as in **Fig. 1**. Scale bars in the merged images, 50 μm. **D-I**. Quantitative analysis of AChR cluster numbers (D), volume (E), number of primary (F) and secondary (G) nerve branches, and the length of primary (H) and secondary (I) nerve branches. Data are expressed as percentage of control (dotted lines). Data are means ± SEM. NS, non-significant, \**p* < *0.05*, \*\**p* < *0.01*, \*\*\**p* < *0.001*, \*\*\*\**p* < *0.0001*; ns, non-significant; ^*#*^*p* < *0.05*, ^*###*^*p* < *0.01; ####p* < *0.0001*: relative to control. *N* = *6* embryos per genotype, one-way ANOVA, Tukey’s or Dunnet’s test.

We next determined the functional relationship between the two proteins during NMJ differentiation *in vivo* in a mouse genetic interaction assay. Two genes are considered to interact genetically if their simultaneous partial knockdown generates a highly penetrant loss-of-function phenotype stronger than that observed following the partial knockdown of either gene alone. We therefore crossed *HSA::Cre;Vangl2*^*LoxP/+*^ and *MuSK*^+/*-*^ mice to generate *MuSK*^+/*-*^;*HSA-Vangl2*^*LoxP/-*^ transheterozygous mice and characterized the resulting NMJ phenotype in diaphragms from E18.5 embryos (**Fig. 6C**). No NMJ differentiation defects relative to control were detected in *HSA::Cre;Vangl2*^*LoxP/+*^ (*HSA-Vangl2*^+/*-*^) and *MuSK*^+/*-*^ heterozygous embryos, except for a slight increase in the length of secondary nerve branches in *MuSK*^+/*-*^ heterozygotes, as previously reported ^14^, and a decrease in the length of primary branches in *HSA-Vangl2*^+/*-*^ mutants (**Fig. 6D-I**). By contrast, *MuSK*^+/*-*^; *HSA-Vangl2*^+/*-*^ transheterozygous embryos displayed profound pre- and postsynaptic abnormalities. The number of AChR clusters was much smaller in *MuSK*^+/*-*^; *HSA-Vangl2*^+/*-*^ than in *HSA-Vangl2*^+/*-*^ and *MuSK*^+/*-*^ embryos (*MuSK*^+/*-*^, 92 ± 7.7 %; *HSA-Vangl2*^+/*-*^, 86 ± 9.0%; *MuSK*^+/*-*^; *HSA-Vangl2*^+/*-*^, 59 ± 8.6%; *p* < 0.001), although their volume was unchanged (**Fig. 6D-E**). The number (*MuSK*^+/*-*^, 85 ± 14.5%; *HSA-Vangl2*^+/*-*^, 124 ± 11.7%; *MuSK*^+/*-*^;*HSA-Vangl2*^+/*-*^, 205 ± 13.1%; *p* < 0.001) and length (*MuSK*^+/*-*^, 148 ± 8.8%; *HSA-Vangl2*^+/*-*^, 108 ± 5.9%; *MuSK*^+/*-*^;*HSA-Vangl2*^+/*-*^, 263 ± 11.5%; *p* < 0.001) of secondary, but not primary nerve branches were significantly increased (**Fig. 6F-I**). *HSA-Vangl2*^*-*/*-*^ and *MuSK*Δ*CRD* ^18^ mouse embryos have similarly altered NMJ phenotypes. We therefore asked whether MuSK/Vangl2 signaling process was mediated by Wnt/MuSK interaction. Similar genetic interaction studies were performed with *HSA::Cre;Vangl2*^*LoxP/+*^ and *MuSK*^*+/ΔCRD*^ mice (**Fig. 6C-I**). As previously reported, *MuSK*^*+/ΔCRD*^ embryos displayed no NMJ formation defects relative to WT embryos ^14^. By contrast, *MuSK*^*+/*Δ*CRD*^;*HSA-Vangl2*^+/*-*^ transheterozygous embryos displayed a partial postsynaptic alteration relative to *HSA-Vangl2*^+/*-*^ and *MuSK*^*+/*Δ*CRD*^ embryos, characterized by a ∼1.5-fold decrease in the volume of AChR clusters (**Fig. 6E**), and presynaptic abnormalities associated with an increase in the number of secondary branches (but not significantly different relative to *HSA-Vangl2*^+/*-*^) and their aimless overgrowth (**Fig. 6H-I**). These results thus revealed an interaction between Vangl2 and MuSK signaling pathways also involving Wnt/MuSK interaction for the regulation of several aspects of NMJ formation.

### Loss of Vangl2 function alters MuSK signaling activity

In a first attempt to characterize the functional role of Vangl2/MuSK interaction in the regulation of AChR clustering, we assessed the impact of the loss of Vangl2 function on MuSK signaling activity (**Fig. 7**). As expected, *Vangl2*^*-*/*-*^ cultured primary myotubes exhibited a strong decrease in Wnt11-induced AChR clustering (**Fig**.**7A and 7B**). Signaling activation mediated by tyrosine kinase receptors is tightly linked to receptor endocytosis^46^. Thus, we next investigated whether AChR clustering deficiency in Vangl2 depleted muscle cells is linked to a defect in Wnt-elicited MuSK signaling activation. *In vitro* Vangl2 knock-down in differentiated muscle cells was achieved by transduction with a lentivirus carrying the shVangl2 sequence, resulting in a decrease in Vangl2 levels by a factor of 10 relative to muscle cells transduced with a scrambled shRNA (shScr), with no effect on MuSK protein levels (**Fig. 7C**). We then monitored the levels of MuSK protein at the plasma membrane in muscle cells transduced with shScr or shVangl2, following Wnt11 stimulation (**Fig. 7D**). Vangl2 depletion did not affect basal levels of MuSK at the plasma membrane. In the shScr cell line, Wnt11 stimulation caused a rapid time-dependent decrease in surface levels of MuSK expression. By contrast, in the ShVangl2 cell line, MuSK surface expression remained unchanged for up to 30 min of Wnt11 stimulation and started to decrease after 1hr of treatment, suggesting that that Vangl2 loss-of-function delayed Wnt11-mediated MuSK internalization. Control experiments showed that the surface levels of TfR expression were unaffected by Wnt11 treatment in both the shScr and shVangl2 cell lines (**Fig. 7D**). Overall, these findings indicate that Vangl2 regulates AChR clustering by controlling the level of Wnt-induced MuSK signaling activity.

**Figure 7.**
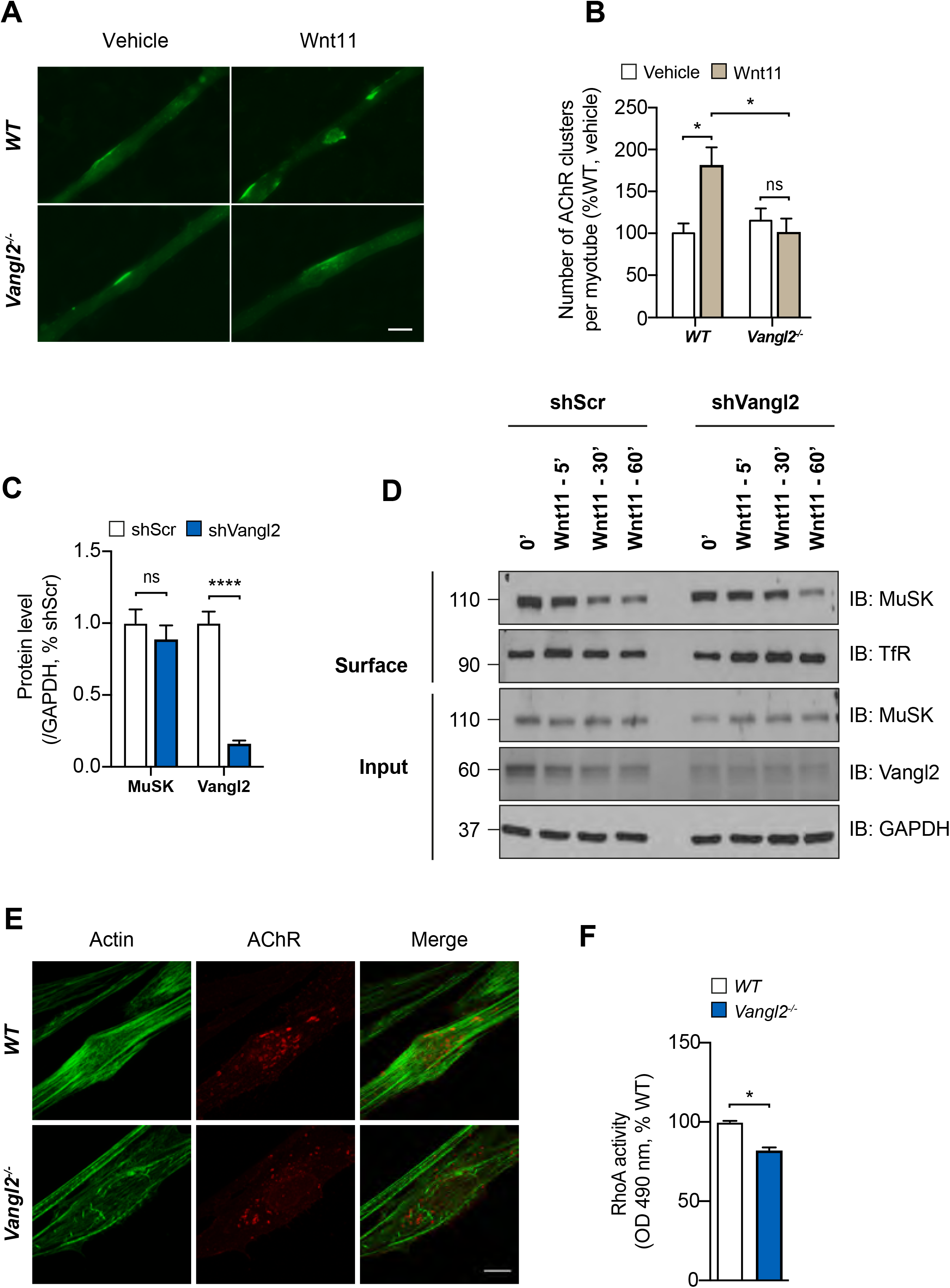
Vangl2 regulates MuSK signaling activity and cytoskeletal organization in muscle cells. **A**. Representative images of WT and *Vangl2-/-* primary myotubes treated or not with Wnt11 recombinant protein and stained for AChR clusters with α-BTX. **B**. Quantitative analysis of the number of AChR clusters. Data are mean ± SEM, ns, non-significant, \**p* < *0.05, N=9* mice per genotype, Mann-Whitney *U* test. **C**. Quantification of MuSK and Vangl2 protein levels normalized to GAPDH in cell lysates from C2C12 myotubes transduced with shScr or shVangl2. Data are means ± SEM. ns, non-significant, \*\*\*\**p* < *0.0001, N* = *6* independent experiments, one-way ANOVA, Tukey’s test. **D**. Western blot analysis of cell surface and total MuSK following Wnt11 stimulation at various time points in C2C12 myotubes transduced with shScr or shVangl2. Transferrin receptor (Tfr), loading control for biotinylated proteins. GAPDH, loading control for input. MW, molecular weight (kDa). IB, Immunoblot. **E**. Representative confocal images of control and *Vangl2*^*-*/*-*^ primary myotubes stained with α-BTX (red, RACh) and Phalloïdin (green, actin). Scale bar in the merged image, 10 µm. **F**. Quantification of RhoA activity in *Vangl2*^*-*/*-*^ myotubes relative to control. Data are means ± SEM, \**p* < *0.05*, Mann-Whitney *U* test, *N* = *3* independent experiments.

Finally, we investigated the impact of loss of Vangl2 function in the regulation of AChR anchoring to the subsynaptic cytoskeleton, to demonstrate further the link between a loss of MuSK signaling activity and AChR clustering deficiency in cells depleted of Vangl2. The protein-protein interaction domains of Vangl2 modulate downstream signaling, including the activation of RhoA GTPases during cytoskeleton organization ^47,48^. We thus characterized differences in the cellular actin cytoskeleton stress fibers arrangement between control and *Vangl2*^*-*/*-*^ myotubes stained with Alexa Fluor 488-conjugated phalloidin to label stress fibers and Alexa Fluor 594-conjugated α-BTX to stain AChR clusters (**Fig. 7E**). Control myotubes exhibited an elongated parallel arrangement of actin stress fibers, which was altered in *Vangl2*^*-*/*-*^ myotubes (**Fig. 7E**). Actin stress fiber arrangement has been shown to be a direct consequence of RhoA activity ^49,50^, and RhoA activity is involved in MuSK-elicited AChR clustering ^51^. Thus, we measured the amount of GTP-bound (active) RhoA in control and *Vangl2*^*-*/*-*^ myotubes. Vangl2-deficient muscle cells had significantly lower levels of RhoA activity than control cells (control, 100 ± 0.7%; *Vangl2*^*-*/*-*^, 82.4 ± 1.5%; *p* < 0.05) (**Fig. 7F**).

## Discussion

This study provides compelling evidence that Vangl2 is part of the core complex of postsynaptic signaling proteins driving the formation and maintenance of neuromuscular synapses throughout life. The constitutive deletion of *Vangl2* confirmed previous findings suggesting a role for Vangl2 in NMJ formation ^15^. However, the pre- and postsynaptic alterations observed in the *Vangl2*^*-*/*-*^ mutant were less severe than the defects previously reported for Vangl2 looptail (Lp) mouse embryos ^15^ indicating a dominant negative function of the Lp mutation on NMJs. This finding is consistent with those of other studies of Vangl2-mediated PCP functions in various cell contexts, showing that Lp mutation leads to the production of a mutant protein interferes with the function of endogenous Vangl2 protein or other central actors of the PCP pathway ^39^.

Our findings reveal uncharacterized functions of Vangl2 in muscle during NMJ formation and maintenance, and notably a cell-autonomous role of Vangl2 in postsynaptic differentiation and NMJ integrity. We found that the conditional deletion of *Vangl2* in muscle altered embryonic NMJ formation, recapitulating the NMJ phenotype of the *Vangl2*^*-*/*-*^ mutant. In contrast, *Vangl2* knockout in motoneurons did not result in NMJ differentiation defects suggesting that muscle Vangl2 cannot be compensated by Vangl2 expression from the motor axons. The muscle-specific deletion of *Vangl2* strongly impaired AChR accumulation in the postsynaptic membrane from the initial nerve-independent step of NMJ formation as highlighted by the small number of aneural and neural AChR clusters, indicating a role for Vangl2 in the formation of immature AChR clusters and in promoting or stabilizing neural AChR clusters. The postsynaptic alterations were accompanied by a sharp overgrowth of motor axons bypassing the remaining AChR clusters and growing all over the muscle surface. These presynaptic deficits were probably caused by the considerable disturbance of the postsynaptic endplates. However, we cannot rule out the possibility that muscle Vangl2 drives the expression of retrograde signaling required for presynaptic motor axon outgrowth ^24^. Several core components of the PCP pathway including Vangl2 are involved in the guidance and/or the outgrowth of a subset of peripheral axons such as commissural, monoaminergic and olfactory axons ^24,25,52–57^. Strikingly, our results show that when Vangl2 expression was ablated specifically in the motoneurons, they reached their final destination and correctly innervated the diaphragm without apparent presynaptic abnormalities, suggesting that motoneuronal Vangl2 is not required cell autonomously for motoneuron axon outgrowth and presynaptic differentiation in the diaphragm muscle. It has been recently shown that the specific innervation of muscle territories by motoneuron pools in the spinal cord requires a subset of core PCP genes. Consistent with this finding, the deletion of *Vangl2*, unlike that of *Celsr3* and *Fzd3*, had no effect on the innervation of hindlimb muscles, indicating that *Vangl2* gene was dispensable for the steering or outgrowth of motor axons in the peripheral nervous system ^53,58^.

Both Vangl2 looptail mutation and *Vangl2*^*-*/*-*^ mice die at birth due to severe neural tube defects, precluding postnatal analyses of Vangl2 function at the NMJ ^59^. In contrast, muscle-specific Vangl2 cKO mutants are viable making it possible to analyze the role of Vangl2 during NMJ maintenance in adult mice. Muscle specific Vangl2 deletion led to NMJ disassembly, as demonstrated by the structural and ultrastructural disorganization of the NMJs including fragmented postsynaptic specializations with reduced junctional folds, sometimes associated with a partial withdrawal of motor axons, resulting in muscle weakness. These structural and functional deficits of NMJ integrity probably result from embryonic defects of NMJ development, but we cannot exclude the possibility that muscle Vangl2 also regulates postnatal NMJ development. Further investigations in mice with inducible Vangl2 mutations, are required to determine the role of muscle Vangl2 more precisely in NMJ maintenance in adult mice.

Our findings reveal that a loss of muscle Vangl2 decreases muscle strength, probably due to an impairment of neuromuscular transmission in both hindlimb and diaphragm muscles in the adult animal. These functional deficits and the dismantling of NMJ structure are similar to the clinical manifestations observed in patients with a congenital myasthenic syndrome (CMS), a genetic neuromuscular disease affecting synaptic transmission at the NMJ ^60,61^. A search of the Rare Disease-connect database ^62^ for CMS-causing genes associated with *Vangl2* gene mutations identified no convincing disease-causing *Vangl2* polymorphisms. Further studies are therefore required. Of note, *Vangl2* mutations are associated with neural tube defects, including spina bifida ^63,64^. Patients suffering from such disorders usually display motor dysfunction in the legs, including the *tibialis anterior* muscle, with mild or severe muscle weakness ^65,66^. This allows to speculate that human *Vangl2* mutations may disrupt Vangl2 function at the NMJ causing the muscle weakness observed in these patients.

So, how muscle Vangl2 regulate AChR clustering? We show here, for the first time, that Vangl2 physically binds MuSK, the master regulator of NMJ development (Burden et al., 2013; DeChiara et al., 1996), and that the two proteins are part of the same genetic pathway *in vivo* suggesting that they act in a common signaling pathway without excluding a possible synergistic effect of two distinct pathways. MuSK/Vangl2 signaling is also partly mediated by Wnt binding to MuSK (via the extracellular CRD of MuSK) consistent with recent data showing that MuSK and Vangl2 have Wnt proteins as common ligands ^15^. Our findings also reveal that Vangl2 controls Wnt-induced MuSK signaling. First, an absence of Vangl2 results in an inhibition of Wnt11-mediated MuSK internalization in muscle cells with no effect on basal MuSK expression at the plasma membrane, indicating a crucial role for Vangl2 in initiating Wnt-induced MuSK endocytosis required for postsynaptic differentiation. Second, Vangl2 depleted muscle cells exhibit reduced Wnt11-induced AChR clustering. Our results are in agreement with Granato’s data in zebrafish showing that Wnt4 and Wnt11r activate MuSK/Unplugged leading to its translocation to recycling endosomes, which is required for the initiation of PCP-like signaling involved in muscle pre-patterning ^19^. Third, the loss of Vangl2 also affected RhoA activity, a well-known downstream effector of MuSK signaling ^51^, leading to a disorganization of the actin cytoskeleton that contributes to AChR clustering defects. These findings demonstrate the critical role of Vangl2 in maintaining MuSK signaling at the level required for its activity to ensure correct postsynaptic differentiation, whilst preventing the widespread, uncontrolled activation of MuSK that probably accounts for the NMJ defects observed in a context of Vangl2 deficiency. The fine-tuning of MuSK signaling activities is crucial to ensure the correct assembly and maintenance of NMJs. MuSK signaling is, therefore, tightly controlled by several processes mediated, for example, by the MuSK adaptor protein DOK7 ^68^. These signaling mechanisms may compensate for Vangl2 deficiency, partly explaining the relatively mild NMJ phenotype and the survival of mice in absence of Vangl2.

Despite the growing evidence of a role for Wnt morphogens in synaptic differentiation in invertebrate and vertebrate organisms ^14,69,70^, the molecular mechanisms underlying Wnt/PCP signaling function in the regulation of NMJ assembly still largely remain elusive. By investigating the role of Vangl2-mediated-Wnt/PCP signaling at the NMJ, we reveal here a more general and crucial function of Vangl2 in regulating MuSK signaling and thereby controlling key aspects of postsynaptic assembly and maintenance, including cytoskeleton organization and AChR clustering.

## Acknowledgments and funding

This work was supported by grants from AFM-Téléthon (grant ID 19546 and 22691) and the Institut National de la Santé et de la Recherche Médicale (INSERM). *In vivo* imaging was performed at the SCM imaging platform of Université de Paris (Plateforme Imagerie du Vivant – PIV), supported by the Ile-de-France Region research program SESAME. We thank Frédérique Rau, Evelyne Bloch-Gallego, Annie Cartaud, Alexandre Dobbertin, Fabien Le Grand and Jean Cartaud for their helpful scientific advices and/or technical assistance. We also thank the core facilities of the Brain and Spine Institute for their contribution to this work (Pheno-ICMice, iGenSeq, ICM-Quant, CELIS and iVECTOR).

## Author contributions

L.S conceived the study. L.S, J.ME, M.B, M.M, N.S and B.F designed the methodology and analyzed the data. M.B, J.ME and L.S wrote the manuscript. M.B performed most of the experiments and analyzed the data. S.C, S.B and C.B contributed to cell culture and biochemical experiments. M.H contributed to confocal image acquisition. A.R, D.L and A.B contributed to electron microscopy analysis. J.MO, M.A, A.F, M.L conducted force measurements and/or electroneuromyography. All the authors read and approved the final manuscript.

## References

1. Li, L., Xiong, W.-C. & Mei, L. Neuromuscular Junction Formation, Aging, and Disorders. Annu. Rev. Physiol. 80, 159–188 (2018).

2. Tintignac, L. A., Brenner, H.-R. & Rüegg, M. A. Mechanisms Regulating Neuromuscular Junction Development and Function and Causes of Muscle Wasting. Physiol. Rev. 95, 809–852 (2015).

3. Barik, A. et al. LRP4 is critical for neuromuscular junction maintenance. J. Neurosci. Off. J. Soc. Neurosci. 34, 13892–13905 (2014).

4. DeChiara, T. M. et al. The receptor tyrosine kinase MuSK is required for neuromuscular junction formation in vivo. Cell 85, 501–512 (1996).

5. Hesser, B. A., Henschel, O. & Witzemann, V. Synapse disassembly and formation of new synapses in postnatal muscle upon conditional inactivation of MuSK. Mol. Cell. Neurosci. 31, 470–480 (2006).

6. Kim, N. et al. Lrp4 is a receptor for Agrin and forms a complex with MuSK. Cell 135, 334–342 (2008).

7. Kim, N. & Burden, S. J. MuSK controls where motor axons grow and form synapses. Nat. Neurosci. 11, 19–27 (2008).

8. Weatherbee, S. D., Anderson, K. V. & Niswander, L. A. LDL-receptor-related protein 4 is crucial for formation of the neuromuscular junction. Dev. Camb. Engl. 133, 4993–5000 (2006).

9. Zhang, B. et al. LRP4 serves as a coreceptor of agrin. Neuron 60, 285–297 (2008).

10. Zhang, W., Coldefy, A.-S., Hubbard, S. R. & Burden, S. J. Agrin binds to the N-terminal region of Lrp4 protein and stimulates association between Lrp4 and the first immunoglobulin-like domain in muscle-specific kinase (MuSK). J. Biol. Chem. 286, 40624–40630 (2011).

11. Gomez, A. M. & Burden, S. J. The extracellular region of Lrp4 is sufficient to mediate neuromuscular synapse formation. Dev. Dyn. Off. Publ. Am. Assoc. Anat. 240, 2626–2633 (2011).

12. Zong, Y. et al. Structural basis of agrin-LRP4-MuSK signaling. Genes Dev. 26, 247–258 (2012).

13. Zong, Y. & Jin, R. Structural mechanisms of the agrin-LRP4-MuSK signaling pathway in neuromuscular junction differentiation. Cell. Mol. Life Sci. CMLS 70, 3077–3088 (2013).

14. Boëx, M. et al. Regulation of mammalian neuromuscular junction formation and maintenance by Wnt signaling. Curr. Opin. Physiol. 4, 88–95 (2018).

15. Messéant, J. et al. Wnt proteins contribute to neuromuscular junction formation through distinct signaling pathways. Dev. Camb. Engl. 144, 1712–1724 (2017).

16. Strochlic, L. et al. Wnt4 participates in the formation of vertebrate neuromuscular junction. PloS One 7, e29976 (2012).

17. Zhang, B. et al. Wnt proteins regulate acetylcholine receptor clustering in muscle cells. Mol. Brain 5, 7 (2012).

18. Messéant, J. et al. MuSK frizzled-like domain is critical for mammalian neuromuscular junction formation and maintenance. J. Neurosci. Off. J. Soc. Neurosci. 35, 4926–4941 (2015).

19. Gordon, L. R., Gribble, K. D., Syrett, C. M. & Granato, M. Initiation of synapse formation by Wnt-induced MuSK endocytosis. Dev. Camb. Engl. 139, 1023–1033 (2012).

20. Henriquez, J. P. et al. Wnt signaling promotes AChR aggregation at the neuromuscular synapse in collaboration with agrin. Proc. Natl. Acad. Sci. U. S. A. 105, 18812–18817 (2008).

21. Jing, L., Lefebvre, J. L., Gordon, L. R. & Granato, M. Wnt signals organize synaptic prepattern and axon guidance through the zebrafish unplugged/MuSK receptor. Neuron 61, 721–733 (2009).

22. Shen, C. et al. Motoneuron Wnts regulate neuromuscular junction development. eLife 7, (2018).

23. Avilés, E. C. & Stoeckli, E. T. Canonical wnt signaling is required for commissural axon guidance. Dev. Neurobiol. 76, 190–208 (2016).

24. Dos-Santos Carvalho, S. et al. Vangl2 acts at the interface between actin and N-cadherin to modulate mammalian neuronal outgrowth. eLife 9, (2020).

25. Shafer, B., Onishi, K., Lo, C., Colakoglu, G. & Zou, Y. Vangl2 promotes Wnt/planar cell polarity-like signaling by antagonizing Dvl1-mediated feedback inhibition in growth cone guidance. Dev. Cell 20, 177–191 (2011).

26. Tissir, F. & Goffinet, A. M. Shaping the nervous system: role of the core planar cell polarity genes. Nat. Rev. Neurosci. 14, 525–535 (2013).

27. Nagaoka, T. et al. The Wnt/planar cell polarity pathway component Vangl2 induces synapse formation through direct control of N-cadherin. Cell Rep. 6, 916–927 (2014).

28. Yoshioka, T., Hagiwara, A., Hida, Y. & Ohtsuka, T. Vangl2, the planar cell polarity protein, is complexed with postsynaptic density protein PSD-95 [corrected]. FEBS Lett. 587, 1453–1459 (2013).

29. Ramsbottom, S. A. et al. Vangl2-regulated polarisation of second heart field-derived cells is required for outflow tract lengthening during cardiac development. PLoS Genet. 10, e1004871 (2014).

30. Wu, H. et al. Distinct roles of muscle and motoneuron LRP4 in neuromuscular junction formation. Neuron 75, 94–107 (2012).

31. Cartaud, A. et al. MuSK is required for anchoring acetylcholinesterase at the neuromuscular junction. J. Cell Biol. 165, 505–515 (2004).

32. Montcouquiol, M. et al. Asymmetric localization of Vangl2 and Fz3 indicate novel mechanisms for planar cell polarity in mammals. J. Neurosci. Off. J. Soc. Neurosci. 26, 5265–5275 (2006).

33. Bolte, S. & Cordelières, F. P. A guided tour into subcellular colocalization analysis in light microscopy. J. Microsc. 224, 213–232 (2006).

34. Jones, R. A. et al. NMJ-morph reveals principal components of synaptic morphology influencing structure-function relationships at the neuromuscular junction. Open Biol. 6, (2016).

35. Sigoillot, S. M., Bourgeois, F., Lambergeon, M., Strochlic, L. & Legay, C. ColQ controls postsynaptic differentiation at the neuromuscular junction. J. Neurosci. Off. J. Soc. Neurosci. 30, 13–23 (2010).

36. Chevessier, F. et al. A mouse model for congenital myasthenic syndrome due to MuSK mutations reveals defects in structure and function of neuromuscular junctions. Hum. Mol. Genet. 17, 3577–3595 (2008).

37. Delacroix, C. et al. Improvement of Dystrophic Muscle Fragility by Short-Term Voluntary Exercise through Activation of Calcineurin Pathway in mdx Mice. Am. J. Pathol. 188, 2662–2673 (2018).

38. Torban, E., Iliescu, A. & Gros, P. An expanding role of Vangl proteins in embryonic development. Curr. Top. Dev. Biol. 101, 237–261 (2012).

39. Yin, H., Copley, C. O., Goodrich, L. V. & Deans, M. R. Comparison of phenotypes between different vangl2 mutants demonstrates dominant effects of the Looptail mutation during hair cell development. PloS One 7, e31988 (2012).

40. Kibar, Z. et al. Ltap, a mammalian homolog of Drosophila Strabismus/Van Gogh, is altered in the mouse neural tube mutant Loop-tail. Nat. Genet. 28, 251–255 (2001).

41. Torban, E., Kor, C. & Gros, P. Van Gogh-like2 (Strabismus) and its role in planar cell polarity and convergent extension in vertebrates. Trends Genet. TIG 20, 570–577 (2004).

42. Arber, S. et al. Requirement for the homeobox gene Hb9 in the consolidation of motor neuron identity. Neuron 23, 659–674 (1999).

43. Brennan, K. J. & Hardeman, E. C. Quantitative analysis of the human alpha-skeletal actin gene in transgenic mice. J. Biol. Chem. 268, 719–725 (1993).

44. Miniou, P. et al. Gene targeting restricted to mouse striated muscle lineage. Nucleic Acids Res. 27, e27 (1999).

45. Thaler, J. et al. Active suppression of interneuron programs within developing motor neurons revealed by analysis of homeodomain factor HB9. Neuron 23, 675–687 (1999).

46. Lemmon, M. A. & Schlessinger, J. Cell signaling by receptor tyrosine kinases. Cell 141, 1117–1134 (2010).

47. Schlessinger, K., Hall, A. & Tolwinski, N. Wnt signaling pathways meet Rho GTPases. Genes Dev. 23, 265–277 (2009).

48. Seifert, J. R. K. & Mlodzik, M. Frizzled/PCP signalling: a conserved mechanism regulating cell polarity and directed motility. Nat. Rev. Genet. 8, 126–138 (2007).

49. Bishop, A. L. & Hall, A. Rho GTPases and their effector proteins. Biochem. J. 348 Pt 2, 241–255 (2000).

50. Zhang, W., Huang, Y. & Gunst, S. J. The small GTPase RhoA regulates the contraction of smooth muscle tissues by catalyzing the assembly of cytoskeletal signaling complexes at membrane adhesion sites. J. Biol. Chem. 287, 33996–34008 (2012).

51. Weston, C. et al. Cooperative regulation by Rac and Rho of agrin-induced acetylcholine receptor clustering in muscle cells. J. Biol. Chem. 278, 6450–6455 (2003).

52. Fenstermaker, A. G. et al. Wnt/planar cell polarity signaling controls the anterior-posterior organization of monoaminergic axons in the brainstem. J. Neurosci. Off. J. Soc. Neurosci. 30, 16053–16064 (2010).

53. Hua, Z. L., Jeon, S., Caterina, M. J. & Nathans, J. Frizzled3 is required for the development of multiple axon tracts in the mouse central nervous system. Proc. Natl. Acad. Sci. U. S. A. 111, E3005–3014 (2014).

54. Lyuksyutova, A. I. et al. Anterior-posterior guidance of commissural axons by Wnt-frizzled signaling. Science 302, 1984–1988 (2003).

55. Tissir, F., Bar, I., Jossin, Y., De Backer, O. & Goffinet, A. M. Protocadherin Celsr3 is crucial in axonal tract development. Nat. Neurosci. 8, 451–457 (2005).

56. Wang, Y., Zhang, J., Mori, S. & Nathans, J. Axonal growth and guidance defects in Frizzled3 knock-out mice: a comparison of diffusion tensor magnetic resonance imaging, neurofilament staining, and genetically directed cell labeling. J. Neurosci. Off. J. Soc. Neurosci. 26, 355–364 (2006).

57. Zhou, L. et al. Early forebrain wiring: genetic dissection using conditional Celsr3 mutant mice. Science 320, 946–949 (2008).

58. Chai, G. et al. Celsr3 is required in motor neurons to steer their axons in the hindlimb. Nat. Neurosci. 17, 1171–1179 (2014).

59. Strong, L. C. & Hollander, W. F. HEREDITARY LOOP-TAIL IN THE HOUSE MOUSEAccompanied by Imperforate Vagina and with Lethal Craniorachischisis When Homozygous. J. Hered. 40, 329–334 (1949).

60. Engel, A. G. Congenital Myasthenic Syndromes in 2018. Curr. Neurol. Neurosci. Rep. 18, 46 (2018).

61. Engel, A. G., Shen, X.-M., Selcen, D. & Sine, S. M. Congenital myasthenic syndromes: pathogenesis, diagnosis, and treatment. Lancet Neurol. 14, 420–434 (2015).

62. Thompson, R. et al. RD-Connect: an integrated platform connecting databases, registries, biobanks and clinical bioinformatics for rare disease research. J. Gen. Intern. Med. 29 Suppl 3, S780–787 (2014).

63. Juriloff, D. M. & Harris, M. J. A consideration of the evidence that genetic defects in planar cell polarity contribute to the etiology of human neural tube defects. Birt. Defects Res. A. Clin. Mol. Teratol. 94, 824–840 (2012).

64. Kibar, Z. et al. Contribution of VANGL2 mutations to isolated neural tube defects. Clin. Genet. 80, 76–82 (2011).

65. Apkon, S. D. et al. Advances in the care of children with spina bifida. Adv. Pediatr. 61, 33–74 (2014).

66. Petronic, I. et al. Distribution of affected muscles and degree of neurogenic lesion in patients with spina bifida. Arch. Med. Sci. AMS 7, 1049–1054 (2011).

67. Burden, S. J., Yumoto, N. & Zhang, W. The role of MuSK in synapse formation and neuromuscular disease. Cold Spring Harb. Perspect. Biol. 5, a009167 (2013).

68. Okada, K. et al. The muscle protein Dok-7 is essential for neuromuscular synaptogenesis. Science 312, 1802–1805 (2006).

69. Klassen, M. P. & Shen, K. Wnt signaling positions neuromuscular connectivity by inhibiting synapse formation in C. elegans. Cell 130, 704–716 (2007).

70. Koles, K. & Budnik, V. Wnt signaling in neuromuscular junction development. Cold Spring Harb. Perspect. Biol. 4, (2012).

