## Supplemental data for "Muscle Van Gogh-like 2 shapes the neuromuscular synapse by regulating MuSK signaling activity"

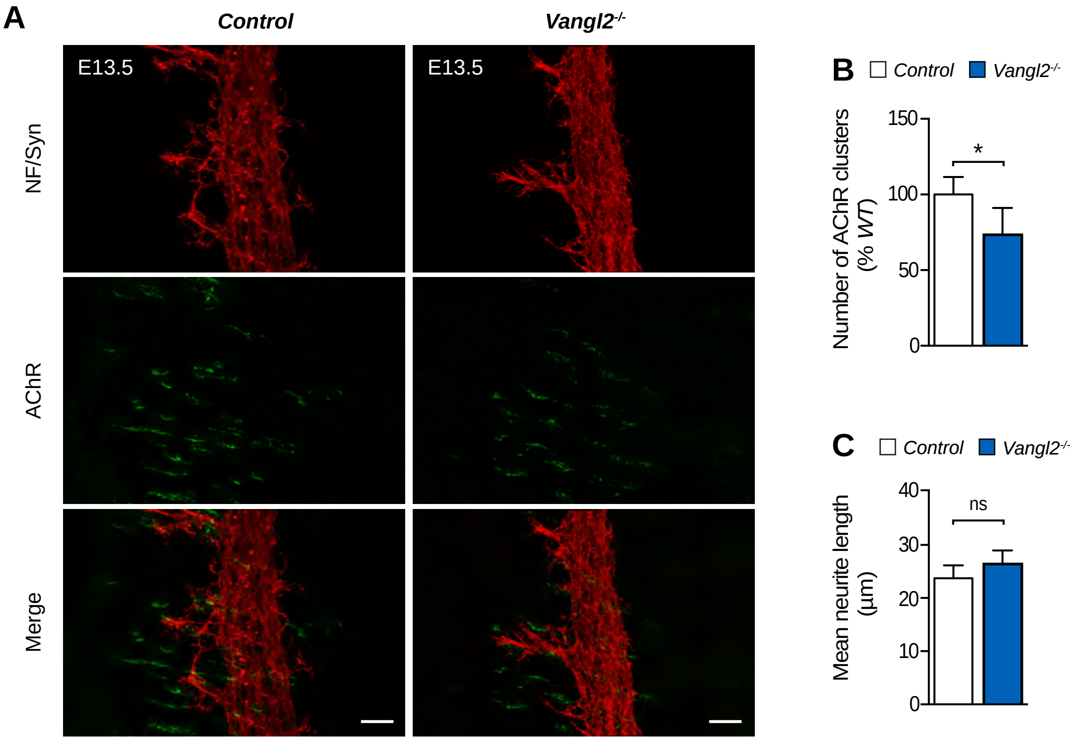

A
Control
Vangl2-/-
E13.5
E13.5
NF/Syn
AChR
Merge
B
Control
Vangl2-/-
Number of AChR clusters (% WT)
150
100
50
0
*
C
Control
Vangl2-/-
Mean neurite length (µm)
40
30
20
10
0
ns

| | | LoxP/+ | | $HSA\text{-}Vangl2^{-/-}$ | |
|---|---|---|---|---|---|
| | | Mean | SEM | Mean | SEM |
| **Phrenic nerve stimulation (indirect stimulation)** | | | | | |
| **Twitch force** | Peak amplitude (mN) | 167.50 | 31.96 | 82.17* | 13.85 |
| | Specific force (mN/mg) | 3.31 | 0.73 | 1.25* | 0.30 |
| **Tetanic force (20Hz)** | Peak amplitude (mN) | 496.20 | 60.25 | 195.90** | 31.91 |
| | Specific force (mN/mg) | 9.69 | 1.99 | 3.50* | 0.95 |
| | Tetanus/twitch ratio | 3.52 | 0.28 | 2.42* | 0.24 |
| **Tetanic force (40Hz)** | Peak amplitude (mN) | 521.40 | 65.88 | 212.70** | 32.01 |
| | Specific force (mN/mg) | 10.36 | 2.36 | 4.25* | 0.97 |
| | Time to peak (ms) | 4.11 | 0.37 | 3.03 | 0.47 |
| **Tetanic force (60Hz)** | Peak amplitude (mN) | 457.00 | 54.28 | 201.30** | 32.88 |
| | Specific force (mN/mg) | 8.92 | 2.02 | 4.02* | 1.05 |
| | Time to peak (ms) | 3.96 | 0.42 | 2.79 | 0.36 |
| **Tetanic force (80Hz)** | Peak amplitude (mN) | 372.40 | 45.17 | 230.00* | 29.31 |
| | Specific force (mN/mg) | 7.14 | 1.56 | 4.85 | 0.86 |
| | Time to peak (ms) | 3.66 | 0.44 | 3.21 | 0.51 |
| **Muscle stimulation (direct stimulation)** | | | | | |
| **Twitch force** | Peak amplitude (mN) | 109.60 | 17.56 | 115.60 | 34.75 |
| | Specific force (mN/mg) | 2.46 | 0.43 | 2.26 | 0.94 |
| **Tetanic force (20Hz)** | Peak amplitude (mN) | 382.70 | 54.15 | 275.20 | 62.65 |
| | Specific force (mN/mg) | 7.66 | 1.72 | 4.46 | 1.40 |
| | Time to peak (ms) | 3.94 | 0.27 | 3.18 | 0.17 |
| **Tetanic force (40Hz)** | Peak amplitude (mN) | 450.40 | 53.89 | 376.40 | 74.72 |
| | Specific force (mN/mg) | 9.06 | 1.90 | 6.94 | 2.25 |
| | Time to peak (ms) | 4.65 | 0.42 | 4.02 | 0.28 |
| **Tetanic force (60Hz)** | Peak amplitude (mN) | 410.30 | 48.65 | 369.60 | 71.52 |
| | Specific force (mN/mg) | 8.32 | 1.79 | 6.89 | 2.21 |
| | Time to peak (ms) | 4.38 | 0.46 | 4.04 | 0.29 |
| **Tetanic force (80Hz)** | Peak amplitude (mN) | 385.40 | 43.67 | 328.70 | 77.19 |
| | Specific force (mN/mg) | 6.59 | 1.15 | 7.29 | 1.98 |
| | Time to peak (ms) | 3.88 | 0.43 | 3.74 | 0.28 |

**Table 1. Muscle tension parameters in *LoxP/+* and $HSA\text{-}Vangl2^{-/-}$ mouse hemidiaphragms.**

**Supplemental Figure 1. Reduced aneural AChR clusters in *Vangl2*$^{-/-}$ mouse embryos**

**A**. Representative confocal images of whole-mount left hemidiaphragms from E13.5 control and *Vangl2*$^{-/-}$ mouse embryos stained with α-BTX (green, AChR), anti-neurofilament (red, NF, phrenic nerve) and anti-synaptophysin antibodies (red, Syn, nerve terminals). Scale bars in the merged images, 20 μm. **B-C.** Quantitative analysis of the number of AChR cluster (B) and mean neurite length (C). Data are means ± SEM, ns, non-significant; $*p < 0.05$, $N = 3$ embryos per genotype, Mann-Whitney *U* test.
